# An Organotypic Oral Squamous Cell Carcinoma Model Recapitulates Epithelial-Stromal Complexity at Single-cell Resolution and Reveals Matrix-derived Signalling as a Therapeutic Target

**DOI:** 10.64898/2026.03.10.710769

**Authors:** Olga S. Yevlashevskaya, Joshua Z. Davies, Jiwon Choi, Shuai Yuan, Arsalan Latif, Gowsihan Poologasundarampillai, Deena M.A. Gendoo, Malgorzata Wiench

## Abstract

Three-dimensional (3D) organotypic cultures recapitulate key structural features of oral tumours and provide controlled, ethical, and reproducible research platforms. However, their ability to faithfully recapitulate in vivo tissue composition and their translational relevance require rigorous validation. Here, we characterised a head and neck squamous cell carcinoma (HNSCC) model using single-cell RNA sequencing to assess maturation, cellular heterogeneity, functionality, and inter– and intra-tissue interactions within epithelial and stromal compartments.

The epithelial layer of the model differentiated into populations closely resembling the tissue of tumour origin, including dividing and precursor cells, heterogeneous basal layer, suprabasal cells, and metabolically specialised populations. All epithelial lineages emerged from proliferative progenitors, driven by dynamic transcription factor programmes. The collagen-rich stroma contained functionally diverse fibroblasts reflecting the heterogeneity of cancer-associated fibroblasts in vivo, including dividing, matrix-producing, immune-responsive, and tumour-like populations. Importantly, extensive epithelial–stromal communication networks developed, essential for cancer epithelium maintenance and tumour microenvironment regulation. Matrix-derived signals, particularly fibronectin, osteopontin, and laminins, constituted dominant inputs to CD44-expressing cancer cells and are associated with patient survival in HNSCC, highlighting potential therapeutic targets.

Overall, this organotypic HNSCC model exhibits high functional fidelity, captures key tumour elements relevant for therapy and resistance and brings confidence in non-animal drug testing.

## 1. Introduction

Head and neck squamous cell carcinoma (HNSCC) is well known for its poor outcomes (37-72% 5-year survival) due to frequent recurrence, metastasis and resistance to therapy (Sung et al., 2021). Only a few targeted therapies have been approved for HNSCC to date, including EGFR inhibitors such as cetuximab and immune checkpoint inhibitors targeting the PD-1/PD-L1 axis (Gougis et al., 2019; Sim et al., 2025). However, meaningful clinical benefit was observed only in a subset of patients, with most cases demonstrating limited survival improvement and low response rates outside selected populations, highlighting the need for alternative therapies. Recent advances in single cell RNA-sequencing and spatial transcriptomics proved key to understanding different cell populations in oral tumours including the supporting role of tumour stroma (Choi et al., 2023; Kürten et al., 2021; Puram et al., 2017; Quah et al., 2023; Xiong et al., 2024). Building on these efforts will require relevant, competent and well-characterised screening platforms that reproduce the complexity identified in vivo. However, the main approaches are either animal models (which diverge from human oral epithelium structure and pathophysiology) or in vitro spheroid cultures (that lack tissue polarity and complexity). With the advent of animal-free and humanised models, organotypic cultures become the most physiologically relevant options characterised by relevant tissue polarity, cell-cell and tumour-stroma interactions, and therefore more accurate drug responses (Macartney et al., 2025; Rodrigues et al., 2021). Importantly, previous HNSCC organotypic models were shown to have expected cancer morphology and represent well patients’ tumours from which they were derived (Gronbach et al., 2020). Nevertheless, current understanding of the cellular and functional complexity, differentiation patterns and cross-tissue interactions in these models remains limited and have not been previously investigated at the single-cell level in oral mucosa cancer organotypic cultures.

HNSCC originates from the squamous epithelium of the mucosa in the oral cavity, larynx and pharynx (Johnson et al., 2020). Complex stratified epithelium forms the outermost layer of mucosal tissues where it creates a homeostatic barrier against abrasion, toxins and infectious agents; its integrity is crucial for maintaining functional healthy tissues. The barrier is established across a multi-layered structure formed by epithelial cells (keratinocytes) and reinforced by 3D cell-cell (desmosomes, tight junctions and adherens junctions) and cell-matrix (hemidesmosomes) interactions (Groeger & Meyle, 2019; Presland & Dale, 2000). In stratified epithelia, the cells form layers representing different stages of maturation, with terminally differentiated cells present on the tissue surface (Fig.1A). The entire structure is constantly replenished by amplifying cells, located either in basal (mouse) or parabasal (human) cell layers (Andl et al., 2016). As the cells migrate towards the surface, they reinforce their interactions and accumulate structural proteins, i.e. cytokeratins (encoded by *KRT* genes). In orthokeratinised epithelium (as seen in gingiva or skin), these processes are clearly visible across spinous, granular and squamous layers. In non-keratinised epithelium (i.e. in buccal mucosa or floor of mouth), the squamous layer is absent, and the maturation is limited to spinous and intermediate layers as shown in Fig. 1A. The epithelial tissue component is separated from the underlying connective tissue by basement membrane, through which it receives nutrients and signals. The connective tissue is composed of ground substance with collagen and elastin fibers, fibronectin, proteoglycans and glycoproteins, primarily produced by fibroblasts.

**Figure 1.**
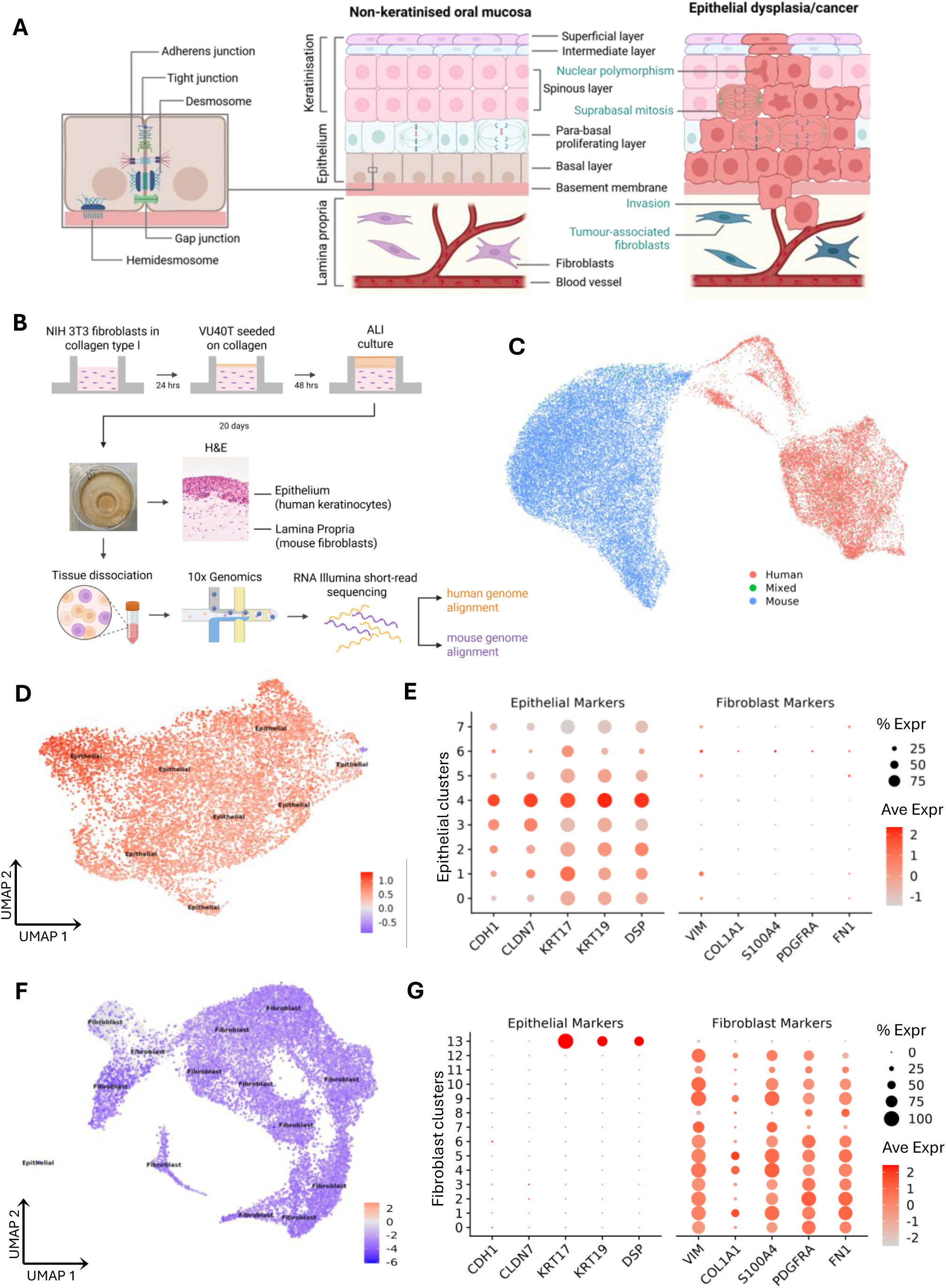
The development of in vitro oral squamous cell carcinoma organotypic model. (**A**) Graphical representation of the oral mucosa structure depicting epithelial and lamina propria components, stratification in non-keratinised epithelium and basement membrane localisation (centre). Left: enlarged section showing cel-cell (desmosomes, tight junctions, adherens and gap junctions) and cell-basement membrane (hemidesmosomes) junctions. Right: oral mucosa changes in oral dysplasia and carcinoma. **(B)** Workflow of the scRNA-seq characterisation of the organotypic oral mucosa model involving: growing OM tissues in vitro using collagen type 1 matrix, NIH/3T3 mouse fibroblasts, VU40T HNSCC keratinocytes and buoyant epithelial culture devices to maintain air-liquid interface (ALI) over 20 days (n=4). This was followed by tissue dissociation, 10X Genomics scRNA-sequencing and alignment to both human and mouse genomes. **(C)** Assignment of single cells to species (human, mouse) allowing for differentiation between epithelial cells and fibroblasts, respectively, for further analysis. The dissociation was achieved by aggregating transcript counts and assigning cell barcodes based on ≥ 70% identity cutoff from each respective alignment. The graph was created by tagging from counting barcodes in both alignments and plotted using human alignment. **(D)** Re-clustering of 9,520 human cells resulting in eight clusters subsequently identified as ‘epithelial’ using manual annotation with predefined markers as per (E). The optimal/most stable number of clusters was determined using clustree at resolution 0.3. **(E)** Dot plot of the common epithelial and fibroblast markers expressed in the human clusters. **(F)** Re-clustering of 22,293 mouse cells resulting in 14 clusters, 13 identified as ‘fibroblast’ and one as ‘epithelial’. The optimal/most stable number of clusters was determined using clustree at resolution 0.6. **(G)** Dot plot of the common epithelial and fibroblast markers expressed in the mouse clusters.

In oral cancer (Fig. 1A, right panel), the epithelial differentiation process is disturbed. This results in unstructured, hyperproliferative tissue, characterised by cellular and nuclear pleomorphism, suprabasal mitotic figures and deviations from normal expression patterns of keratins and junctional proteins (Groeger & Meyle, 2019; Rivera & Venegas, 2014). In invasive cancer, malignant cells acquire the ability to invade the lamina propria, which is facilitated by changes in tumour microenvironment. In response to signals from cancer and immune cells, fibroblasts in the underlying lamina propria become activated and are referred to as cancer associated fibroblasts (CAFs) (Chhabra & Weeraratna, 2023; D. Yang et al., 2023). CAFs secrete growth factors, cytokines and chemokines that influence cancer cell behaviour and tumour growth and modify extracellular matrix (ECM) to support cancer cell spreading (Custódio et al., 2020). Although tumour microenvironment is a well-recognised contributor to carcinogenesis, CAFs are still poorly understood due to their heterogeneity and dynamic behaviour (Cords et al., 2023; D. Yang et al., 2023).

Key aspects of HNSCC biology including differentiation, signalling, barrier function, drug metabolism, cancer-stroma interactions, can only be reliably studied using 3D models that ascertain physiological context, architecture and microenvironment (Macartney et al., 2025; Rodrigues et al., 2021). These epithelial rafts involve layers of epithelial cells with increasing levels of differentiation growing on top of fibroblast-containing matrix, most commonly collagen hydrogel, recapitulating the structure presented in Fig.1A. The differentiation process requires constant maintenance of the epithelial cell layer at the air-liquid interface (ALI), where the epithelial cells are exposed to air while LP remains in contact with cell culture medium (Hewitt et al., 2022).

In this study, we establish a reproducible organotypic oral cancer model and, using single-cell transcriptomics and immunostaining, investigate whether it recapitulates cellular and functional complexity that is critical of HNSCC tumorigenesis. The analysis validates the functional fidelity of the model against the existing in vivo data. Additionally, it uncovers previously underappreciated cellular states and alternative maturation programmes in stratified tumour epithelium. It also demonstrates the feasibility of fibroblast cell line in vitro divergence into multiple relevant CAF subtypes and maps the cross-tissue communication network. Finally, it identifies matrix-derived signalling as potential therapeutic target.

## 2. Results

### 2.1. Single-cell RNA-seq of in vitro oral mucosa cancer tissues developed as dual-lineage, dual-species 3D model

Epithelial organotypic rafts were generated using human HNSCC human papillomavirus (HPV)-negative cell line VU40T, which originated from base of tongue tumour, and immortalised mouse embryonic NIH 3T3 fibroblasts embedded in type 1 rat tail collagen to mimic lamina propria. First, collagen gels with fibroblasts were established in buoyant epithelial culture devices (Hewitt et al., 2022). After 24 hours, the VU40T cells were seeded on top of the gels and allowed to form monolayers, before being lifted to air liquid interface and cultured over 20 days to form stratified epithelia (Fig. 1B). Four single cell libraries were prepared using 10X Genomics Chromium and sequenced. Due to the presence of human and mouse cells in the model setup, the reads were aligned to both genomes for respective analysis of epithelial cells and fibroblasts. The replicates showed very high similarity (Suppl. Fig. 1A-C) and QC metrics (Suppl. Fig. 1D-E); therefore, the reads were combined, and the integrated data were used in subsequent analyses and visualisation using the uniform manifold approximation and projection (UMAP) plots.

The initial analysis revealed that each genome alignment resulted in two distinct clusters separating epithelial and mesenchymal lineage features (Suppl. Fig. 2A-B). It was suspected that conserved sequences combined with short read sequencing allowed for permissive inter-species alignment. To avoid cross-species contamination between the two tissue types in subsequent genome analyses, the data from both genome alignments were combined and each cell was classed as ‘human’ or ‘mouse’ based on ≥ 70% reads identity cutoff with a small number of cells showing ‘mixed’ identity (Fig. 1C, Suppl. Fig. 2C). The resulting 9,520 human-tagged cells and 22,293 mouse-tagged cells were re-clustered to characterise the epithelial and lamina propria components of the model, respectively.

Following the re-clustering, all ‘human’ clusters had epithelial identity (Fig. 1D) and consistently expressed main epithelial cell-specific markers (E-cadherin (*CDH1)*, claudin (*CLDN7)*, keratins *KRT17* and *KRT19*, desmoplakin (*DSP)*) while lacking fibroblast-specific markers (vimentin (*VIM)*, collagen *COL1A1*, *S100A4*, platelet-derived growth factor receptor alpha (*PDGFRA)*, fibronectin (*FN1*)) (Fig. 1E). Similarly, the majority (13 out of 14) of ‘mouse’ clusters were identified as fibroblasts with exception of one small cluster of epithelial cells (cluster 13), which was excluded from further analysis (Fig. 1F). This was confirmed by the fibroblast clusters expressing at least two classic fibroblast markers and entirely lacking the epithelial markers (Fig. 1G). The re-clustered human and mouse data were used to characterise the epithelial and lamina propria components of the model, respectively.

### 2.2. Epithelial component of the model consists of cell populations relevant to both oral mucosa and squamous cell carcinoma biology

Following the initial analysis of the scRNA-seq data, the cells identified as of human origin (n=9,520) were re-clustered at clustree resolution 0.3 (Suppl. Fig. 3). This resulted in eight epithelial clusters, with cluster 0 accounting for a quarter, and cluster 7 for only 1.6% of all epithelial cells (Fig. 2A). The clusters formed a connected structure rather than discrete clouds, indicating gradual change in cellular identities across the tissue. To characterise the contribution of each cluster in the structure of the formed stratified epithelium, the expression of keratins, components of cell-cell junction (adhesion junctions, desmosomes, tight junctions) and cell-matrix junctions (hemidesmosomes and basement membrane proteins) was investigated (Fig. 2B) followed by pathway analysis conducted using REACTOME database (Fig. 2C). Cluster 4 (12% of all cells) showed the highest enrichment in genes encoding for keratins (including *KRT5*, *KRT19* and *KRT6A*) and all cell-cell junctional proteins, including components of desmosomes (desmocollins (*DSCs*), desmogleins (*DSGs*) and plakoglobin (*JUP))*, tight junctions (claudins (*CLDNs*) and tight junction proteins *TJP1/TJP2*) as well as adherens junctions (cadherins (*CDH*), vinculin (*VCL*) and nectin) (Fig. 2B, D), indicating these cells are part of suprabasal, differentiating epithelial layers (Cluster E-Suprabasal). This was further evident in enrichment of pathways, such as keratinisation and cell-cell junction organisation (Fig. 2C). By comparison, the expression of genes involved in formation of basement membrane and hemidesmosome attachments was lower in cluster 4 when compared to clusters 1, 2, 5 and 6. The most superficial layer also appears to be involved in immune signalling (Fig. 2C) and shows the highest enrichment for genes encoding innate immune response proteins including interleukins (Suppl. Fig. 4A).

**Figure 2.**
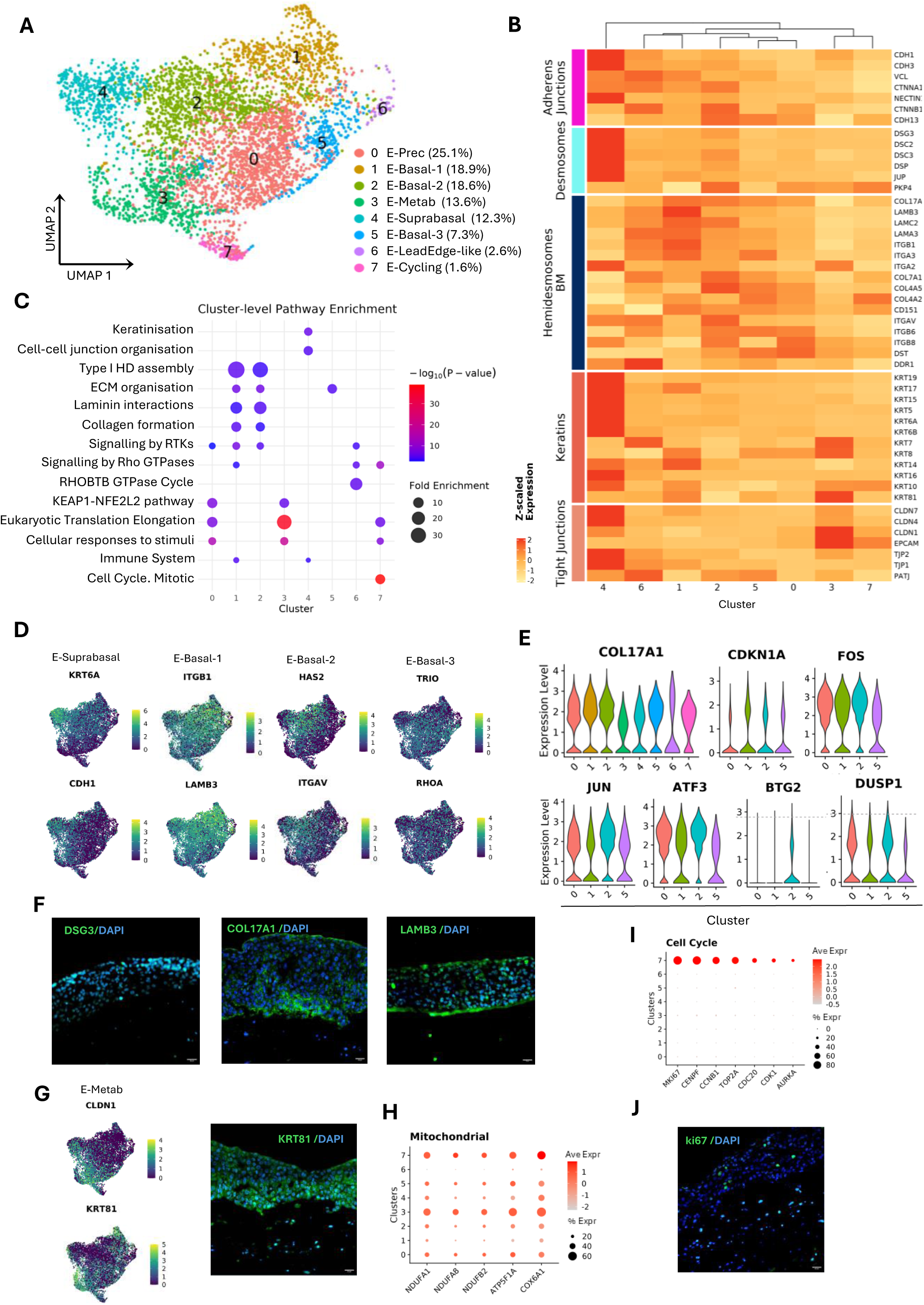
Characterisation of the epithelial component of the organotypic model. (**A**) Seurat clustering of epithelial cells using uniform manifold approximation and projection (UMAP). The clusters are colour-coded, and their labels and percentage tissue contribution are indicated in the legend. **(B)** Heatmap showing expression of genes associated with keratinisation, cell-cell and cell-basement membrane epithelial junctions z-scaled across the epithelial clusters and ordered by hierarchical clustering. **(C)** Pathway enrichment analysis performed using REACTOME database and based on the top 200 markers from each cluster. To create the graph top 10 pathways for each cluster with FDR <0.05 were combined and manually created to avoid redundancies. Dot size indicates fold enrichment with colour-coded p-value. Some GO terms are truncated for brevity; original names, pathways ID and full gene lists are provided in Suppl. File 2. **(D)** Feature plots showing RNA expression of selected markers for clusters E-Suprabasal, E-Basal-1, E-Basal-2 and E-Basal3. **(E)** Violin plots of relative gene expression for genes associated with previously described non-proliferative basal epidermal states (Haensel et al., 2020): *COL17A1*+, early response (*FOS*, *JUN*, *ATF3*, *BTG2*, *DUSP1*) and growth arrested (*CDKN1A*). **(F)** Immunostaining of the markers for suprabasal (DSG3) and basal/basement membrane (COL17A1, LAMB3) epithelial layers in organotypic models at day 20. Counterstained with DAPI. Scale bars, 20 µm. **(G)** Feature plots showing RNA expression of structural protein markers enriched in cluster E-Metab (left) and immunostaining for one of them, KRT81 (right). Scale bar, 20 µm. **(H)** Gene expression dot plot for genes associated with mitochondrial activity in the epithelial clusters. **(I)** Gene expression dot plot for genes associated with cell cycle and proliferation in the epithelial clusters. **(J)** Immunostaining for proliferation marker Ki67. Scale bar, 20 µm.

Clusters 1, 2 and 5 include diverse groups of basal cells, all characterised by high expression of *COL17A1*, laminins and integrins (Fig. 2B), and are annotated as E-Basal-1, E-Basal-2 and E-Basal-3, respectively. All basal clusters show involvement in ECM organisation; in addition, E-Basal-1 and E-Basal-2 are enriched in pathways leading to collagen formation and hemidesmosome assembly (Fig. 2C). Cluster E-Basal-1 is particularly enriched in expression of laminins (*LAMB3*, *LAMC2*, *LAMA3*) and integrins *ITGB1* and *ITGA3* (Fig. 2B, D), indicating direct contact with basement membrane. By comparison, cluster E-Basal-2 is not only enriched in cell-ECM structures (including collagens type 4 and 7, and *ITGAV)* but also have high expression of components of desmosomes and adherens junctions, sharing the cell-cell attachment function with E-Suprabasal cluster 4. The top marker for E-Basal-2 is *HAS2*, gene encoding hyaluronan synthase 2, crucial for ECM formation, adhesion, cell motility and tumour metastasis (Passi et al., 2019) (Fig. 2D). Cluster E-Basal-3 differs from E-Basal-1 and –2 by enrichment in Rho-GTPases pathways. These signalling pathways become dominant in cluster 6 (Fig. 2C), primarily due to expression of genes involved in dynamic reorganisation of actin cytoskeleton (e.g. *RHOA*, *ROCK2*, *TRIO,* (Fig. 2D)) and collagen-activated RTK (*DDR1*) (Fig. 2B). In epithelial cells, activation of Rho-GTPases leads to a switch from keratin intermediate filament to actin-based cytoskeleton and associated intercellular and cell-matrix junctions (Parri, 2010). The cell population in cluster 6 indeed shows a decrease in the expression levels of keratins and keratin-associated desmosomal elements while retaining their adhesion junctions, laminins and integrins (Fig. 2B). In cancer epithelium such switch is associated with migratory properties and invasiveness, while in wound healing – with the epithelial leading edge (Infante & Etienne-Manneville, 2022), hence the cluster annotation is E-LeadEdge-like.

In addition, cluster 0 is also enriched in some of the elements of basement membrane attachments, including *COL17A1* (Fig. 2B, E). As this is the most abundant cluster and does not appear to be committed to any function in stratified epithelium, it is considered here a precursor cluster (E-Prec). The observed basal clusters show distinct similarities to the non-proliferative basal states previously described in mouse epidermis: Col17a1+, Fos+ early response state (ER) and Cdkn1a+ growth arrested state (GA) (Haensel et al., 2020). Although all basal clusters (0, 1, 2, 5, 6) show high expression of *COL17A1*, E-Basal-2 cluster displays features of the reported ER subcluster through enrichment for inflammatory response (*NFKB1* program discussed later) and consistent expression of *FOS*, *JUN*, *ATF3*, *BTG2* and *DUSP1* (Fig. 2E). Cluster E-Basal-1 is *CDKN1A*-positive and shows lowest enrichment for metabolic activity, oxidative phosphorylation and oxidative stress response (shown later), reminiscent of the GA subcluster.

Therefore, the established oral cancer mucosa model shows differentiating suprabasal population enriched in cell-cell junctions and heterogenous basal populations. As expected, E-Suprabasal cells are present on the surface of the epithelium (shown by DSG3 immunostaining) and above the COL17A1-positive cells from basal clusters while LAMB3 overlaps with the basement membrane, consistent with the location of E-Basal-1 cluster (Fig. 2F).

Cluster 3 is characterised by a unique transcription profile with high expression of tight junction components, especially *CLDN1* (Fig. 2B, G) and comparatively low enrichment for desmosomes or hemidesmosomes (Fig. 2B). The expression of keratins is unique and shows an opposing pattern to E-Suprabasal cluster with high transcription of *KRT81*, *KRT7* and *KRT8* (Fig. 2B), while immunostaining also shows more intermediate location of the cells (Fig. 2G). This cluster is described here as Cluster E-Metab due to high metabolic activity evidenced by increased translation (Fig. 2C) and mitochondrial activity (Fig. 2H) leading to oxidative stress and activation of oxidative stress response (KEAP1-NFE2L2 pathway, Fig. 2C). Cluster 0 is also characterised by oxidative stress response but without the tight junctions’ enrichment and with moderate expression of keratins, components of adhesion junctions and hemidesmosomes.

None of the clusters described so far shows evidence of cell proliferation, therefore they are either post-mitotic/differentiating or quiescent. The only cluster enriched for cell cycle and proliferation markers is cluster 7 (E-Cycling) with high expression of *MKI67*, *TOP2A*, *CENPF*, *CCNB1* amongst others (Fig. 2I). As expected for proliferating cells, E-Cycling cluster is characterised by increased metabolism and mitochondrial function (Fig. 2C, H). Although these cells express both keratins and cell-cell and cell-matrix junctions’ components, the expression is low compared to other clusters. The Ki67 immunostaining shows these cells are sparsely scattered in para– and supra-basal layers (Fig. 2J).

The identified cell populations show clearly that once the HNSCC established cell line was placed in 3D environment with signals provided from connective tissue and air liquid interface, it recapitulated in vivo epithelial cancer tissue, including amplifying cells, precursor cells, heterogenous basal layer, suprabasal layer and metabolic cells.

### 2.3. Epithelium maturation is guided by changes in transcription factors’ expression

To further understand the functional tissue development, maturation events and accompanying changes in transcriptional regulation were investigated. Epithelial maturation trajectory was explored using Monocle3 (Trapnell et al., 2014). The pseudotime paths were overlayed with the epithelial component UMAP (Fig. 3A) and the distance from the proliferating E-Cycling cluster 7 cells was calculated for all the remaining clusters (Fig. 3B). It confirmed that most cells transitioned into precursor cells (E-Prec cluster 0) before differentiating into cells connected to basement membrane and ECM (clusters E-Basal-1-3 (clusters 1, 2 and 5). These cell populations are heterogenous as they separate at the exit from E-Prec cluster. Cluster E-Basal-1 becomes the most differentiated cluster committed to interactions with basal lamina. Cells in cluster E-Basal-3 move away from classical epithelial features and become more dependent on actin cytoskeleton and associated signalling through Rho-GTPases. This trajectory continues into cluster 6. On the other hand, the E-Suprabasal cluster 4 differentiates from the cells of E-Basal-2 cluster with increase in cell-cell junctions and keratins expression, the characteristics already observed in E-Basal-2 (Fig. 2B). Therefore, E-Suprabasal and E-Basal-1 are the most differentiated cell populations for suprabasal and basal trajectory, respectively (Fig. 3A-B). E-Metab cluster 3 appears to develop independently of basal/suprabasal pattern and directly from E-Cycling cluster 7 cell populations, confirming the previously noted uniqueness of the cluster.

**Figure 3.**
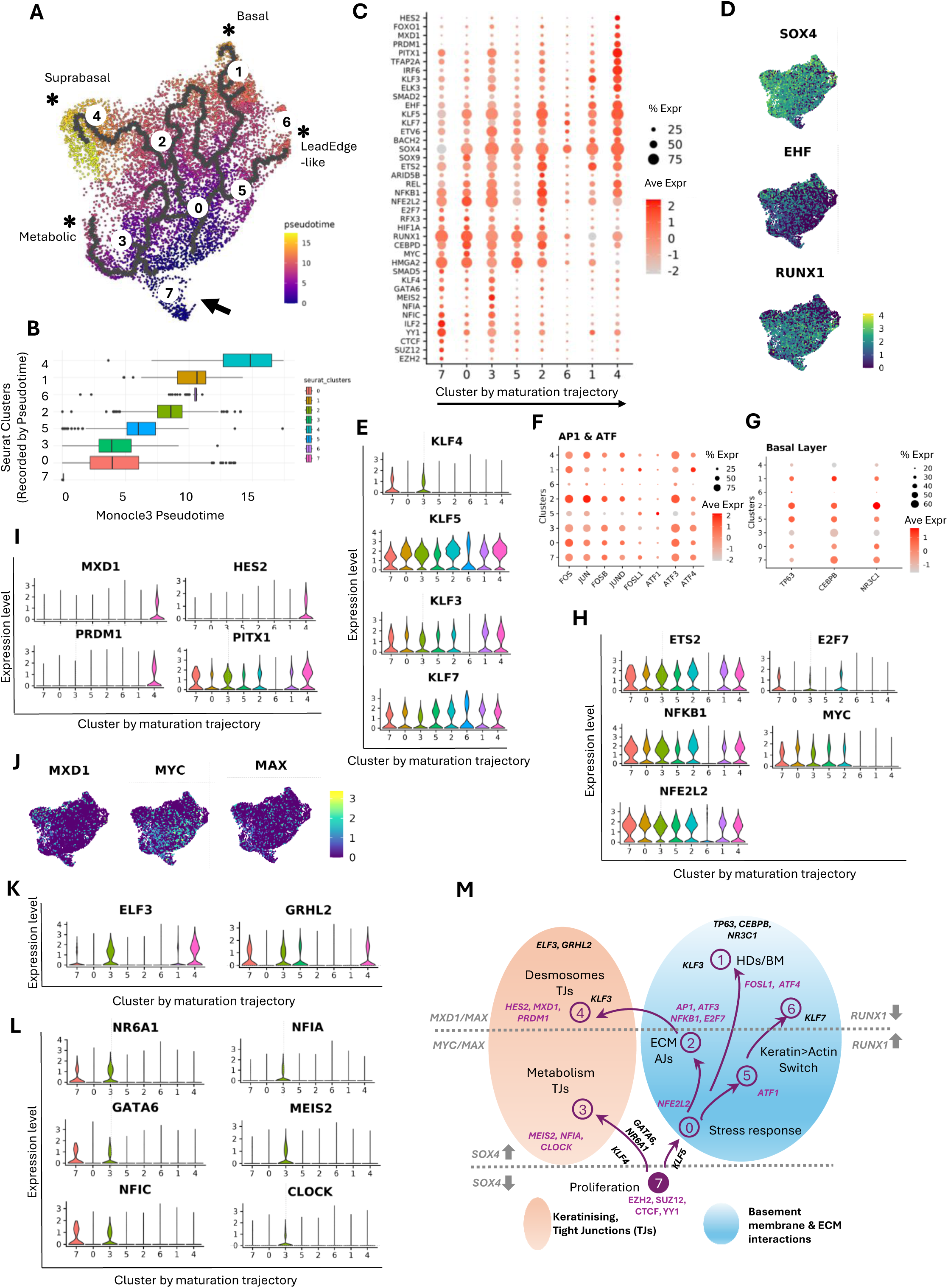
In vitro epithelial maturation and the associated changes in transcription factors’ expression. (**A**) Pseudotime inference of epithelial populations shown in UMAP plot. The arrow indicates proliferating cells in cluster 7 (E-Cycling). Asterisks show end points of the four main differentiation trajectories. **(B)** Epithelial differentiation progression calculated by Monocle 3 pseudotime distance from E-Cycling cluster 7. **(C)** Dot plot of gene expression changes for selected transcription factors that follow the maturation pseudotime trajectory. The clusters are shown in the order of maturation distance. **(D)** Feature plots depicting RNA expression of *SOX4*, *EHF* and *RUNX1* and their changes across maturation trajectory. **(E)** Violin plots of relative gene expression for selected KLF transcription factors. Note the cluster order follows the maturation trajectory. **(F-G)** Gene expression dot plots for (**F**) transcription factors from AP-1 and ATF families and (**G**) transcription factors associated with basal epithelial layers (clusters 1, 2 and 5 (E-Basal-1-3)). **(H)** Violin plots of relative gene expression for selected transcription factors associated with main expression pathways in E-Basal-2 (cluster 2) and E-Prec (cluster 0). **(I)** Violin plots of relative gene expression for transcription factors associated with suprabasal differentiation culminating at E-Suprabasal cluster 4. **(J)** Feature plots of maturation-dependent expression changes of genes encoding for components of MYC/MAX and MXD1/MAX complexes. **(K-L)** Violin plots of relative gene expression of transcription factors associated with (**K**) epithelial clusters enriched in tight junctions (E-Metab cluster 3 and E-Suprabasal cluster 4) and (**L**) with metabolic activity in cluster 3. **(M)** Graphical representation of maturation changes during in vitro epithelium development towards basal cell and tight junction/keratinising –related phenotypes. The associated transcription factors are indicated for clusters and trajectories. The dotted lines show the major switch in the indicated TFs expression as related to the level of maturation. TJs, tight junctions; AJs, adherens junctions.

Tissue maturation is known to follow changes in gene expression patterns, controlled by transcription factors (TFs). Therefore, the expression of TFs in the model was next investigated, and the most apparent pattern that emerged reflected the differentiation trajectory and transition from amplifying cells to precursor cells and finally basal/suprabasal populations (Fig. 3C). E-Cycling cluster is associated with expression of progenitor cell markers *EZH2* and *SUZ12* as well as *CTCF*, *YY1* and *ILF2* TFs, and the transition from amplifying to differentiating cells is marked by the switched-on expression of *SOX4*, absent in the proliferating cells (Fig. 3C-D). *RUNX1* expression marks the proliferative and early, less-differentiated, epithelial cells and decreases in the most differentiated clusters, whereas *EHF* expression increases gradually with maturation (Fig. 3C-D). This is expected as EHF is an epithelium-specific (Ets family) TF, key in establishing epithelial differentiation and barrier function (Luk et al., 2018). These dynamics indicate a transition from priming factors (RUNX1) (Deltcheva & Nimmo, 2017) to plasticity modulators (SOX4) (Moreno, 2019) and, ultimately, to epithelial establishing factors (EHF), consolidating epithelial barrier programs.

KLF (Krüppel-like) TFs, particularly KLF4 and KLF5, are crucial for regulating epithelial differentiation and homeostasis and their expression is often dysregulated in squamous cell carcinomas in a context-dependent manner (Li et al., 2015; McConnell et al., 2007). In the in vitro model, low *KLF4* expression is only present in E-Cycling and E-Metab clusters, while high levels of *KLF5, KLF7* and *KLF3* are widespread (Fig. 3E). Both down-regulation of *KLF4* and up-regulation of *KLF5* and KLF7 are associated with dedifferentiation in HNSCC, promoting tumour progression and invasiveness (Cai et al., 2024; Li et al., 2015; Yang et al., 2021). On the other hand, in the described model, *KLF3* expression is highest in the most differentiated clusters 1 and 4, suggesting a role in epidermal differentiation (Fig. 3E).

Activator Protein 1 (AP-1) and Activating Transcription Factors (ATFs) are also well known in regulation of epithelial biology (Eckert et al., 2013; Wang et al., 2021). AP-1 functions as a dimer of proteins from FOS (FOS, FOSB, FOSL1, FOSL2) and JUN (JUN, JUNB, JUND) families. Although the AP-1 genes, as well as closely related *ATF3* and *ATF4* factors are widely expressed across the clusters, E-Basal-2 is the most enriched in majority of them apart from *FOSL1,* which is highly expressed in E-Basal-1 (Fig. 3F).

Independent of the maturation stage, all clusters with basal features have high expression of *TP63*, *CEBPB*, and *NR3C1* (Fig. 3G). *TP63* is a master regulator essential for the formation of stratified epithelia, also involved in squamous cell carcinogenesis (Koster & Roop, 2004; Moses et al., 2019). Its expression is specific to the basal cell layer for which it serves as a marker (Koster & Roop, 2007). The direct association of basal epithelium with *NR3C1* and *CEBPB h*as not been previously described. In addition to E-Basal-2 being enriched in the above-mentioned TFs, E-Basal-2 is also characterised by high expression of several TFs known to be involved in both stratified epithelia formation/maintenance and in HNSCC development: including *NFKB1*, *MYC*, *E2F7* and *ETS2* (Fig. 3H). Cluster 0 is primarily involved in response to stimuli and the KEAP-NFE2L2 pathway. In agreement with this, it shows the highest expression of *NFE2L2* transcription factor (Fig. 3H). *NFE2L2*, which encodes for NRF2, plays key role in protecting oral mucosa from oxidative stress by activating antioxidant and cytoprotective genes (Wakamori et al., 2022). In summary, the cells associated with basal phenotype are regulated through TP63, while several HNSCC-associated and oxidative stress-related TFs are highly expressed in E-Basal-2 and E-Prec, respectively.

The E-suprabasal cluster possesses the most differentiated characteristics within the model and several transcription factors are indeed associated with them (Fig. 3I): PITX1 is a developmental TF involved in craniofacial patterning and epithelial maturation of oral mucosa (Overmiller et al., 2024); *HES2* expression in the suprabasal layer indicates transient delay in differentiation to ensure the stratification (Kageyama et al., 2007); *PRDM1* drives terminal differentiation of epithelial cells (Magnúsdóttir et al., 2007) and this is further supported by *MXD1* up-regulation. MXD1 competes with MYC for binding to their partner MAX and by doing so acts as a repressor and antagonises pro-proliferative MYC activity (Hurlin & Huang, 2006). The mutually exclusive expression of *MYC* and *MXD1* is clearly seen between less and more differentiated clusters (Fig. 3J). Altogether, it points towards E-Suprabasal cluster 4 as post-mitotic cells (*MXD1*), committed to terminal differentiation (*PRDM1* and *PITX1*) but still at a pre-terminal state (*HES2*).

Despite the differences between clusters 3 (E-Metab) and 4 (E-Suprabasal), they also share common enrichment for tight junctions (Fig. 2B). In agreement with it, these two clusters are the most enriched for *ELF3* and *GRHL2* (Fig. 3K), which are epithelial-specific TFs whose expression indicates polarised, barrier-tight epithelium. GRHL2 is a master regulator of epithelial polarity, tight junctions and apical-basal barrier integrity (Tanimizu & Mitaka, 2013) while ELF3 enforces epithelial identity and supports TJs assembly and maintenance (Kohno et al., 2006). E-Metab stems directly from proliferating cells and, together with TJs, shows high metabolic and mitochondrial activity. This is reflected on the regulatory level by cluster-specific expression of *NFIA*, *CLOCK* and *MEIS2*, as well as *KLF4*, *NR6A1*, *GRHL2*, *GATA6* and *NFIC* with expression restricted to clusters 3 and 7 (E-Cycling) (Fig. 3L). It likely points at trajectory commitment towards mature transporting barrier where KLF4 marks epithelial commitment towards epithelial lineage and, together with GRHL2/ELF3, promotes epithelial polarity and TJ program. This could lead to high membrane turnover demands, provided for by NR6A1-driven lipid metabolism (Wang et al., 2019) and NFIA-linked oxidative phosphorylation (Hiraike et al., 2023).

Merging the maturation trajectory with expression of transcription factors shows the development of two programmes: keratinising, TJ-rich epithelia and BM/ECM-associated epithelia (Fig. 3M). The cells associated with basal phenotype are regulated through TP63, while several HNSCC-associated and oxidative stress-related TFs highly expressed in E-Basal-2 and E-Prec, respectively. Tight junctions are established via two independent events: either directly from the cycling cells and maintaining *KLF4* expression (accompanied by metabolic/secretory programme) or following a classic epithelial keratinisation trajectory guided by *MXD1*/*PRDM1*/*HES2*. These changes overlay the general differentiation-dependent pattern of *RUNX1*, *SOX4* and *EHF* expression.

### 2.4. The in vitro model features majority of oral cancer therapeutic targets and therapy resistance mechanisms

So far, the single cell reconstruction of the in vitro model showed a well-developed, although not terminally differentiated epithelium, in many structural aspects mimicking in vivo tissue with clear epithelial differentiation trajectories driven by cancer-like changes in transcription factors’ expression. The epithelial raft culture of VU40T cells resulted in stratified epithelium typically composed of around 15-20 layers of cells (Fig. 4A). The basal cell layer appeared to be tightly attached to both lamina propria and the neighbouring cells with some clusters of cells invaginating into the lamina propria and detaching from the basal layer, indicating possible invasive properties. By comparison, the cells in para-basal layers, forming the largest component of the tissue, are large and more loosely attached to each other, with no distinct strata as expected in cancer epithelium. These transition into layers of smaller cells characterised by stronger attachments and eventually exhibiting high level of keratinisation at the surface, shown by the eosin staining. The immunostaining for DSG3, COL17A1, LAMB3 and KRT81 (Fig. 2F, 2G) confirmed the suprabasal location of cluster 4, basal/para-basal locations of clusters 1, 2, 5 and 0 and cluster 3 location in the para-basal/middle epithelial layers.

**Figure 4.**
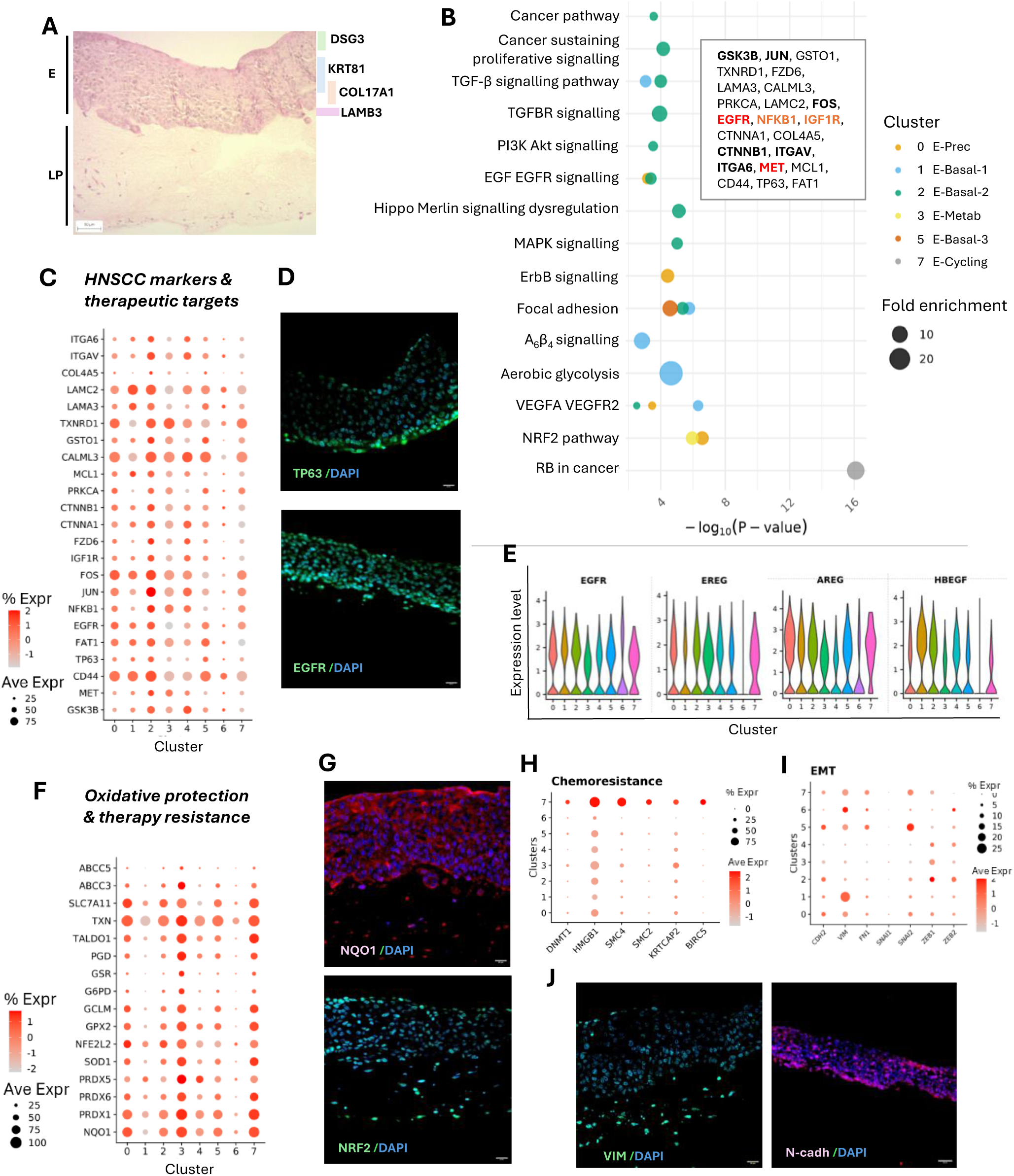
HNSCC characteristics of the in vitro model. (**A**) Hematoxylin and eosin (H&E) staining of representative organotypic model following 20 days of ALI culture (E: epithelium, LP: lamina propria). Scale bar, 50 µm. The approximate localisation of staining for DSG3, KRT81, COL17A1 and LAMB3 is based on the images shown in Figure 2. **(B)** Pathway enrichment based on top 200 markers for each cluster and Wikipathways database. The pathways with FDR <0.05 from each cluster were combined, manually curated and reduced based on the similarity and gene list overlap. The graph shows the selected pathways’ enrichment in epithelial clusters. The enriched clusters are colour-coded as shown in the legend. X axis indicates p-value, dot size – fold enrichment. Some GO terms are truncated for brevity; original names, pathways ID and full gene lists are provided in Suppl. File 3. The inset includes HNSCC-related therapeutic targets involved in the indicated pathways and present in the model; approved/emerging in red/bold and potential in bold. Clusters 4 and 6 are not included as they did not show enrichment in any of the selected pathways. **(C)** Gene expression dot plot in the epithelial clusters for genes reported as HNSCC markers and therapeutic targets. **(D)** Immunostaining of p63 and EGFR in the organotypic model at day 20, counterstained with DAPI. Scale bars, 20 µm. **(E)** Violin plots of relative gene expression for genes encoding EGFR and its ligands. **(F-G)** Oxidative stress response in the epithelial component of the organotypic model. (**F**) Gene expression dot plot for genes involved in cyclooxygenase and thioredoxin reductase systems and (G) immunostaining of the main transcription factor NRF2 and direct responder NQO1. Counterstained with DAPI. Scale bars, 20 µm. **(H)** Gene expression dot plot for genes reported previously to be associated with HNSCC chemoresistance via removal of cell cycle checkpoints. **(I-J)** Expression of EMT markers in the organotypic model. (**I**) Gene expression dot plot for EMT-related genes (note the scaled % Expr legend). (**J**) Immunostaining of VIM and N-cadh in the organotypic model at day 20. Counterstained with DAPI. Scale bars, 20 µm.

Next, we aimed to establish the relevance of the tissue mimics for HNSCC research, both for studying biological mechanisms and in determining therapeutic targets and mechanisms underlying chemo– and radiotherapy resistance. Several pathways are known to be implicated in HNSCC carcinogenesis: TGF-β, PI3K/Akt, EGF/EGFR, MAPK, ErbB, Hippo, JAK/STAT, Wnt/β-catenin and Notch (Constantin et al., 2024). The epithelial clusters were investigated for the enrichment in these pathways. Cluster E-Basal-2 stands out as being associated with majority of them (Fig. 4B) in addition to ‘Cancer pathways’, ‘Hallmark of cancer sustaining proliferative signalling’ and ‘Head and neck squamous cell carcinoma’. This is primarily due to expression of genes such as *GSK3B*, *MET*, *NFKB1*, *EGFR*, *FOS, JUN, IGF1R*, *FZD6*, *PRKCA*, *CTNNA1*, *CTNNB1*, *COL4A5*, *FAT1*, *CD44* and *TP63* (Fig. 4B-C). Many of these genes are therapeutic targets in HNSCC: established/approved (*EGFR*, *MET*), emerging (*IGF1R*, *NFKB1*) or potential (*GSK3B*, *JUN*, *FOS*, *ITGAV*, *ITGA6*, *CTNNB1*). The immunostaining confirms the basal P63 expression while EGFR expression extends above this layer (Fig. 4D).

Of particular interest is the approved EGFR-targeting therapy. The EGF/EGFR and/or ErbB signalling is prominent in E-Basal-2 (cluster 2) and E-Prec (cluster 0) which express not only *EGFR* but also EGFR ligands *AREG, EREG* and HBEGF (Fig. 4E). Furthermore, in the described model, the top marker for E-Basal-2 is *HAS2* (encoding Hyaluronan Synthase 2, Fig. 2D) that produces large glycoprotein hyaluronan (HA). HA promotes EGFR oncogenic signalling by forming complex with CD44 (HA-CD44) which then interacts with EGFR (Wang & Bourguignon, 2006) whereas EGFR signalling itself upregulates *HAS2* expression (Pienimäki et al., 2001). All elements of this positive feedback loop that leads to chemotherapy resistance are highly expressed in E-Basal-2 cells. In addition, the resistance can be established by activation of alternative signalling pathway involving insulin growth factor receptor (IGF1R) (van der Veeken et al., 2009), which is also highly expressed in the model. Interestingly, CD44 surface marker which is most commonly used to isolate cancer stem cells (CSC) in HNSCC (Prince et al., 2007), appears to be widely expressed in our model similarly to other HNSCC CSC markers (Suppl. Fig. 4B).

A common mechanism underlying radio– and chemotherapy resistance involves cytoprotection offered by activated oxidative stress response pathway and its master regulator NFE2L2 (NRF2) (Sporn & Liby, 2012). E-Metab cluster 3, E-Prec cluster 0 and E-Cycling, which together account for ∼40% of cells, show strong enrichment in KEAP1-NFE2L2 pathway (Fig. 2C) together with upregulation of oxidative stress response genes in both cyclooxygenase and thioredoxin reductase (TXN) systems (Fig. 4F). Down-regulation of these pathways in HNSCC cells results in exacerbated oxidative stress and cancer cell death (Gleneadie et al., 2021). NRF2 and NQO1 immunostaining confirms widespread expression of both markers of oxidative stress response, nuclear and cytoplasmic, respectively (Fig. 4G).

Resistance to cancer therapies can also be conferred via removing the blocks on cell cycle checkpoints and apoptosis. In the model, the E-Cycling cluster is uniquely associated with a panel of genes previously linked to multidrug chemoresistance in HNSCC tumours (Khera et al., 2023): *HMGB1, SMC4*, *SMC2*, *BIRK5*, *AURKA* and *DNMT1* (Fig. 4H).

The main prognostic marker in cancer treatment is the ability to invade locally and spread to distant locations. For tumours originating from epithelium, this process involves epithelial-mesenchymal transition (EMT), a phenotype switch during which the cells loose intercellular epithelial junctions and acquire mesenchymal markers (CDH2/N-cadherin, VIM/vimentin) and motility. It is a complex progress directed by a set of TFs: SNAI1, SNAI2, TWIST, ZEB1 and ZEB2 (Nieto et al., 2016). These classical markers, associated with single cell EMT are absent or very weakly expressed in the epithelial clusters (Fig. 4I), indicating the single cell EMT is likely absent in the model. However, 10-20% of cells in E-Basal-3 express *CDH2*, *FN1* and *SNAI2* while a small fraction of cells in E-LeadEdge-like strongly express *VIM* and *ZEB2*. In addition, around 25% of E-Basal-1 cells have moderate *VIM* expression. Both vimentin and N-cadherin are primarily detected by immunostaining in the basal layers (Fig. 4J). Together with activation of Rho-ROCK and actin pathways in E-Basal-3 and E-LeadEdge-like, coupled with *DDR1* (collagen-activated RTK) expression, it could support collective migration along collagen fibres and partial EMT (p-EMT, (Parri & Chiarugi, 2010)).

In summary, the in vitro model represents squamous epithelium with clear activation of HNSCC-driving pathways, expression of cancer markers and therapeutic targets, as well as the presence of cytoprotective mechanisms and potentially p-EMT. This makes it a valuable platform for testing cancer therapies.

### 2.5. Lamina propria of the model contains heterogenous CAF populations derived from dividing fibroblasts and forming complex tumour stroma environment

Fibroblasts are the main cell type responsible for formation and homeostasis of connective tissues, including lamina propria. Therefore, in vitro lamina propria populated with fibroblasts supports the growing sheet of epithelial cells and allows for epithelial polarity and paracrine signalling. In cancer, CAFs help to create stromal environment and secrete growth factors, cytokines and chemokines that influence cancer cell behaviour, tumour growth and invasion (Chhabra & Weeraratna, 2023; Lendahl et al., 2022; Sahai et al., 2020). Both normal fibroblasts and CAFs are very plastic and heterogenous (Bensa et al., 2023 Cords et al., 2023; Lendahl et al., 2022). An increasing number of scRNA-seq studies address this heterogeneity and characterise transcriptionally distinct groups residing in tumour microenvironment. The two main fibroblast groups that consistently emerge across studies are (i) fibroblast responsible for ECM deposition and remodelling, and (ii) pro-inflammatory fibroblasts characterised by production of cytokines and anti-microbial peptides and interactions with T cells and macrophages (Yang et al., 2023). In relation to CAFs they are often referred to as myCAFs (myofibroblastic CAFs) or mCAFs (matrix-producing), and iCAFs (inflammatory or immunomodulating). In addition, other types have been described and included: meCAFs (metabolic CAFs, characterised by metabolite secretion, translation, glycolysis and gluconeogenesis), tumour-like tCAFs, interferon-response CAFs (ifnCAFs) and dCAFs (dividing CAFs) (Bensa et al., 2023; Cords et al., 2023; Mou et al., 2023).

In this study, we used NIH/3T3 fibroblasts, an immortalised embryonic mouse cell line. During the epithelial raft culture, the cells established themselves in collagenous matrix and matured into populations of different functions. After the initial species and fibroblast assignment, 21,737 cells were confirmed to be of mouse origin and were further re-clustered at clustree resolution 0.6 (Suppl. Fig. 5) which resulted in 13 clusters (Fib-0 to Fib-12) (Fig. 5A). To characterise them, the expression of the typical fibroblast markers (*Fn1*, *Col1a1*, *Col1a2*, *S100a4*, Vim, *Pdgfra*, *Dcn*, *Sparc*) (Bensa et al., 2023; Lendahl et al., 2022) was first assessed (Fig. 5B). Collagens are the primary fibre forming proteins in lamina propria. Fibronectin is an adhesive glycoprotein present in ECM, which binds to collagen and integrin and is important in myofibroblast differentiation (Belhabib et al., 2021). Vimentin forms intermediate filaments of fibroblast cytoskeleton, while Pdgfra (platelet-derived growth factor receptor) is a tyrosine kinase receptor involved in signalling, key for fibroblast proliferation and survival as well as myofibroblast transition. S100a4, also known as fibroblast specific protein 1 (Fsp1) plays a role in fibroblast activation and acts both intracellularly and in the ECM, triggering pro-inflammatory responses and ECM remodelling (Belhabib et al., 2021). Genes encoding for fibronectin (*Fn1*), vimentin (*Vim*) and *S100a4* were the most commonly and strongly expressed in most clusters, with cluster Fib-7 lacking *Fn1* and *Pdgfra*, and cluster 8 not expressing *Vim* and *S100a4*. Clusters Fib-7 and –8 are strikingly deprived of synthesis of collagens, which are primarily expressed in clusters 1, 4, 5, 9 and 12 (Fig. 5B). *Sparc* has similar expression to *Col1a1* and *Col1a2*, while *Dcn* expression extends to clusters 2, 3 and 7 (Fig. 5B). Dcn (proteoglycan decorin) and Sparc (glycoprotein also known as osteonectin or basement membrane protein 40) are both ECM components that bind to collagen and facilitate fibres assembly and maturation (Belhabib et al., 2021; Zhang et al., 2018).

**Figure 5.**
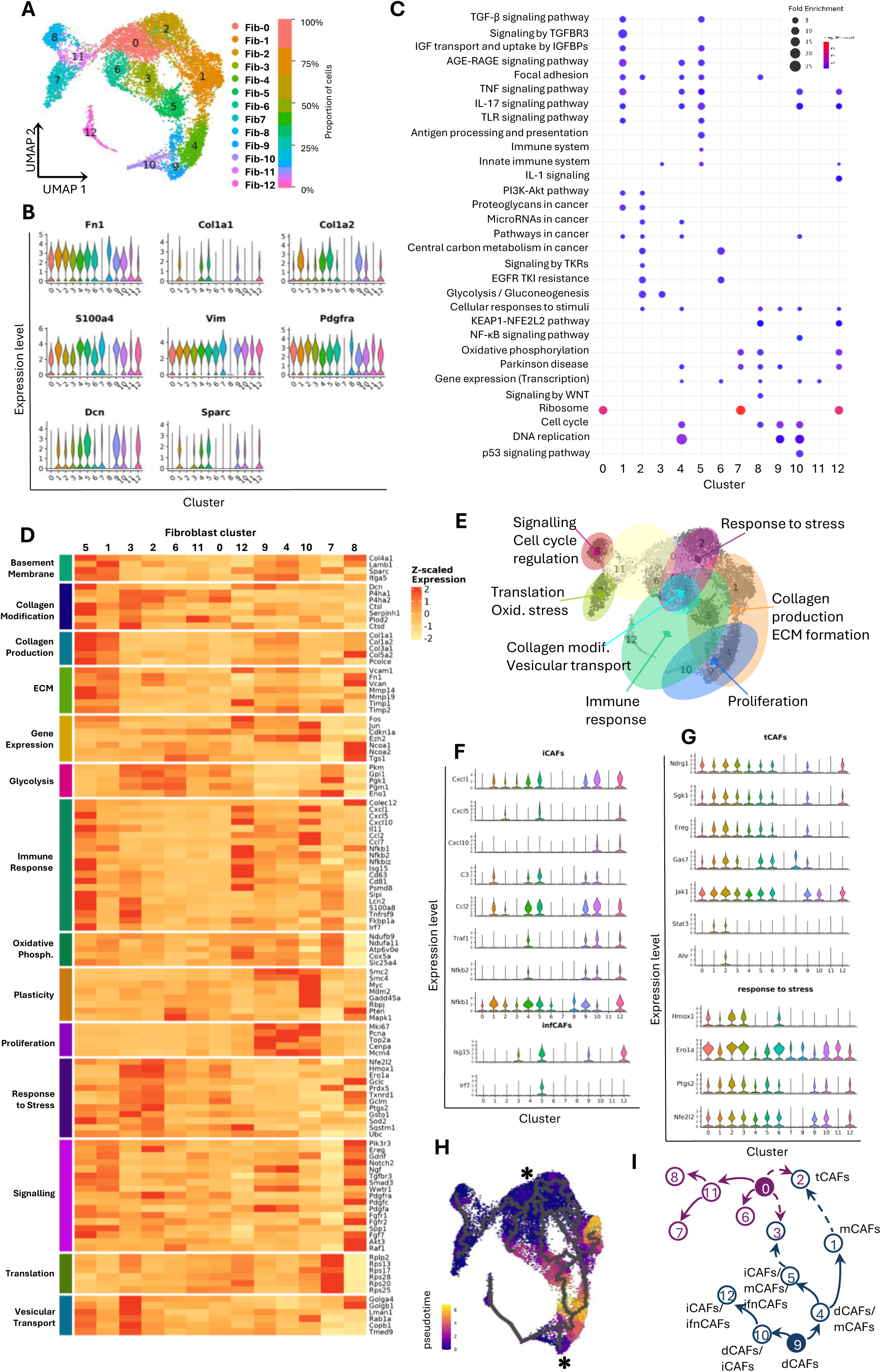
Characterisation of lamina propria component of the organotypic model. (**A**) UMAP representation of Seurat clustering of fibroblast populations. The clusters are colour-coded and their labels and percentage tissue contribution are shown to the right. **(B)** Violin plots of relative gene expression for common fibroblast markers in all fibroblast populations. **(C)** Pathway enrichment analysis based on the top 200 markers from each cluster and combined KEGG/Wikipathway/REACTOME databases. The pathways were selected to best represent he fibroblast heterogeneity. Dot size indicates fold enrichment with colour-coded p-values. Some GO terms are truncated for brevity; original names, pathways ID and full gene lists are provided in Suppl. File 5. **(D-E)**. Functional characterisation of the fibroblast clusters. (**D**) Heatmap showing expression of genes associated with cell functions which were selected based on pathway enrichment analysis and guided the (**E**) general overview of overlapping properties of the fibroblast clusters. **(F-G)** Violin plots of relative expression of genes previously associated with (**F**) inflammatory CAFs (iCAFs) and interferon-response CAFs (ifnCAFs), and (**G**) tumour-like CAFs (tCAFs) enriched for cell response to stress/stimuli genes. **(H-I)** Origins and differentiation of the fibroblast populations in the organotypic model. (**H**) Pseudotime inference of fibroblast populations shown in UMAP plot. The two source populations are indicated by asterisks. (**I**) Schematic overview of the trajectories with fibroblast clusters’ numbers and CAFs types indicated.

To further characterise the fibroblast subtypes, pathway enrichment was assessed using combined KEGG, REACTOME and WIKIPATHWAYS databases (Fig. 5C). The representative genes were selected from the most enriched and significant groups and manually curated to show the diverse fibroblast functions (Fig. 5D). High expression of genes involved in matrix synthesis (including collagens and fibronectin) and remodelling (matrix metalloproteinases (*Mmp*) and tissue inhibitors of metalloproteinases (*Timp*)) was evident in clusters Fib-5, and Fib-1 and to a lesser extent in Fib-9 and Fib-4. Fib-3 mostly expressed genes involved in collagen modifications (i.e. *P4ha1*, *Ctsd*), which was coupled with strong enrichment for vesicular transport and response to stress. This indicates that populations in Fib-1 and Fib-5 are primarily involved in ECM formation regulated by an active TGF-β pathway (Fig. 5C), while cells from cluster 3 potentially secrete proteins supporting collagen fibres formation in the matrix. In addition, cluster Fib-5 expressed genes involved in basement membrane formation (indicating potential vicinity) and was enriched in most genes from the immune system category. Clusters Fib-12 and Fib-10 expressed cytokines, chemokines and nuclear factor kappa B (*Nfkb*) genes, also reflected by enrichment for NF-κB pathway. Genes involved in response to stress (primarily oxidative) were most expressed in Fib-2 and Fib-3. Clusters 9, 4 and 10 all expressed *Mki67*, *Pcna* and *Top2a* genes, indicating dividing cells (Fig. 5D; Suppl. Fig. 6); however, they differed by the contribution to matrix formation (Fib-4 and Fib-9 but not Fib-10) and immune response (only Fib-10). In addition, Fib-10 highly expressed *Smc4*, *Myc*, *Gadd45a* and *Rbpj* indicating high plasticity.

In summary, the analysis performed so far shows highly heterogeneous fibroblast groups with partially overlapping functionalities (Fig. 5E), representing the previously described CAF populations: dCAFs (Fib-9, Fib-4, Fib-10; ∼15% of cells), mCAFs (Fib-1, Fib-5 and to some extent Fib-4; ∼30% of cells) and iCAFs (Fib-5, Fib-10 and Fib-12; ∼12% of cells) (Fig. 5F). A distinct interferon response-enriched fibroblast subtype has been previously highlighted in bladder cancer (Liu et al., 2025) and breast cancer (Cords et al., 2023) and primarily characterised by expression of *ISG15*. In our model, high expression of *Isg15* was detected in Fib-5 and Fib-12, which overlapped with *Irf7* expression in Fib-5 (Fig. 5F). Therefore, ifnCAFs populations have also developed in the in vitro model, with (Fib-5) and without (Fib-12) mCAFs component, similarly to F-POSTN and F-ISG15 populations described by Liu et al. (2025).

tCAFs have been previously characterised by gene expression pattern similar to tumour cells, with upregulation of genes involved in cancer pathways, metastasis and migration (Cords et al., 2023). One of the main markers for this population is *NDRG1*, which has the highest expression in Fib-2, Fib-3 and Fib-0 in our data set (Fig. 5G). In addition, Fib-2 has the strongest expression of *Sgk1*, *Ereg*, *Gas7*, *Jak1*, *Stat3*, and *Ahr* (Fig. 5G) linked to enrichment for cancer and PI3K/Akt pathways (Fig. 5C), high energy consumption (glycolysis) and activated response to stress (Fig. 5G), suggesting this subpopulation is the equivalent of tCAFs in the in vitro model.

Fibroblast populations in the upper-left part of the UMAP exhibit low involvement in the typical matrix-producing and inflammatory fibroblast functions, with clusters 0 and 11 lacking distinctive characteristics, while Fib-6 showing enrichment for glycolysis, and Fib-7 and Fib-8 developing translational and transcriptional/signalling activities, respectively. The development of these clusters can be potentially explained by the time trajectory analysis. The pseudotime mapping revealed two origins with starting points at Fib-9 and Fib-0 (Fig. 5H). Fib-9 consists of dCAFs, which transition to two more dCAF populations, specialising in matrix production (Fib-4, dCAFs/mCAFs) and inflammatory responses (Fib-10, dCAFs/iCAFs) (Fig. 5I). Fib-10 further progresses towards very distinct iCAFs/ifnCAFs (Fib-12) and Fib-4 towards large population of Fib-1 (mCAFs) and to Fib-5 with features of both mCAFs and iCAFs, including ifnCAFs. The second starting node is placed within Fib-0 which is the largest cluster accounting for 17% of all fibroblasts. It is possible this cluster represents the originally seeded fibroblasts, which, although established in the matrix, do not proliferate to give rise to daughter cells further able to develop the main CAFs functions. However, through transitional Fib-11, the cells form Fib-7 and Fib-8 populations. It is not clear from the pseudotime trajectory whether Fib-2 tCAFs develop from the original cell population or follow the trajectory from the dCAFs via cluster Fib-1.

### 2.6. The model develops complex epithelium-stromal interactions which highlight matrix proteins-derived signalling as the main support for cancer cells and potential therapeutic target

In the described in vitro model, cancer epithelium and CAFs-containing lamina propria form accurate representations of their in vivo equivalents. As such, they are expected to maintain a supportive crosstalk. Here, we used the CellChat (Jin et al., 2021) to investigate the intra– and inter-tissue signalling. The incoming signals clearly separated into two patterns: one received by epithelial cells and the other by fibroblasts (Fig. 6A). The main signal receiver in the epithelium is E-Basal-2 (cluster 2), which represents the cells enriched for cancer pathways. Within the fibroblasts’ populations, signals are primarily received by Fib-1, Fib-4, Fib-5 and Fib-9 (Fig. 6A). These clusters are also predominantly involved in producing the outgoing signals. The outgoing signalling can be separated into three patterns (Fig. 6B). All outgoing fibroblast signalling is represented by pattern 2. Outgoing pattern 1 originates from the less differentiated epithelial cells (clusters 0, 2, 5, 7) and accounts for majority of the signalling pathways. It differs from the signalling produced by the more differentiated epithelial clusters (1, 4, 3, 6), which is represented by direct cell-cell contacts (desmosomes, cadherins, JAM proteins, claudins) in addition to IGFBP and PLAU pathways (Suppl. Fig. 7).

**Figure 6.**
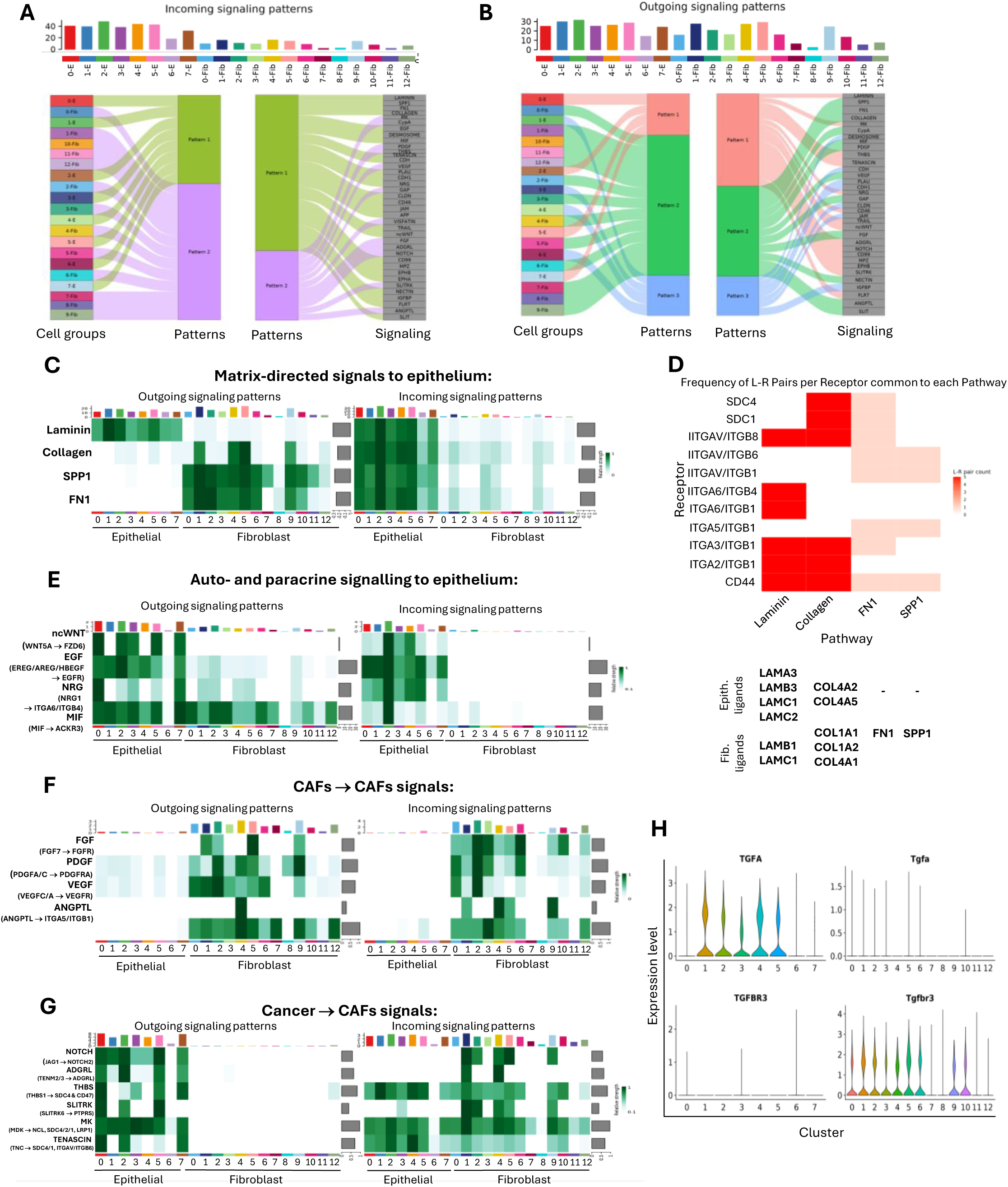
Intra– and inter-tissue cross talk developed in the organotypic model. (**A-B**) Alluvial plots showing incoming (**A**) and outgoing (**B**) signalling for each epithelial and fibroblast cluster grouped into patterns determined by CellChat (left-side graphs). The signalling pathways involved in each pattern are indicated on the right-side graphs. The contribution of each cluster to the incoming and outcoming communication is indicated above the alluvial plots via bar graphs. **(C-D)** Signalling from the components of ECM received by cancer epithelium cells. (**C**) Heatmap with relative importance of each cell cluster in producing ligands (outgoing signalling) and receiving the signals via associated receptors (incoming signalling). (**D**) Interdependent expression of the specific receptors and receptor dimers in epithelial cells and associated ligands, based on the computed network centrality measures for each of the signalling pathways: laminins, collagens, FN1 and SPP1. **(E-G)** Relative contribution of each cell cluster in the incoming and outgoing signals for the indicated pathways involved in auto– and paracrine signalling (**E**) received by epithelial cells, (**F**) signalling between CAFs and (**G**) signalling received by CAFs via ligands produced by epithelial cells. The main ligands and receptors involved are indicated for each signalling pathway. **(H)** Violin plots of relative gene expression of the TGF-β pathway factors expressed by epithelial and fibroblast components of the model: ligand *TGFA* and receptor *TGFBR3*.

Epithelium receives signals both from the matrix interactions and via auto/paracrine ligands. The majority of the matrix-generated signals originate from lamina propria components: collagen, fibronectin and osteopontin (encoded for by *SPP1*) produced by mCAFs (Fig. 6C). In addition, E-Basal-1 (cluster 1) produces laminins that interact with other epithelial clusters. Independent of the inducer, the signals are received by cancer cells via similar receptors: integrins (ITGAV, ITGA6, ITGA3, ITGA2, ITGB1, ITGB4, ITGAB8), CD44 and syndecans (SDC4, SDC1) (Fig. 6D, Suppl. Fig. 8A). The strongest interactions are observed between CD44-FN1 (Suppl. Fig. 8B), CD44-SPP1 (Suppl. Fig. 8C), CD44-LAMB3, CD44-LAMC2, CD44-LAMA3 and ITGA2B1 with LAMC2 and LAMB3 (Suppl. Fig, 8D).

The auto/paracrine signals received by epithelial cells include WNT5A (ncWNT pathway), EGFR ligands (EREG, AREG, HBEGF), NRG1 and MIF (Fig. 6E). The ncWNT signalling is entirely intra-epithelial with FZD6 as main receptor, while EGF and NRG ligands, although primarily epithelial, are also backed by their supply from CAFs. E-Basal-2 cluster is the main signalling receiver for both matrix-guided and paracrine signals. This is most striking in MIF signalling where all epithelial and stromal cells supply the ligand for the benefit of E-Basal-2 cells via ACKR3 receptor (Fig. 6E). The unique E-Metab cluster 3 maintains its specific signalling by responding to ephrin (EPHA/B pathways) and TNF signalling through Amyloid Precursor Protein (APP) and TNF-Related Apoptosis Inducing Ligand (TRAIL) signals from epithelium. It also exchanges signals with the stroma by responding to SLIT signalling and producing NAMPT (VISFATIN pathways) (Suppl. Fig. 9).

Similarly, CAFs are influenced by both fibroblast and epithelial signals. FGF, PDGF and VEGF signalling, well known in supporting CAFs’ functions, in addition to the less established ANGPTL and CypA, are almost entirely contained within the stroma (Fig. 6F), suggesting the presence of self-sustaining pro-proliferative, pro-survival, and angiogenic loops. On the other hand, JAG1, TENMT2-3 and SLITRK6 ligands are only produced by the epithelial cells to be received only by CAFs via NOTCH2, ADGRL and PTPRS receptors, respectively (Fig. 6G). In addition, ligands produced in the epithelium activate THBS, MK/MDK and Tenascin signalling in both fibroblasts and epithelial cells (Fig. 6G). The cancer-to-stroma signals are produced by less differentiated epithelial clusters (0, 2, 5 and 7) and received primarily by dividing fibroblasts (Fib-1, –4, –5, –9) and by t-CAFs (Fib-2). The quiescent fibroblasts in Fib-0, Fib-11, Fib-7 and Fib-8 are least represented in signalling pathways. Similarly, the iCAFs clusters Fib-10 and Fib-12 are missing from the recognised pathways. Since they are shown to express chemo– and cytokines (Fig. 5), this is likely due to the missing partners of the immune component, which is absent in the model. Although the master fibroblast regulator, TGF-β signalling, was not identified in the CellChat analysis, it was represented in the model by *TGFA* ligand expression in most epithelial clusters and *TGFBR3* receptor expression in fibroblasts (Fig. 6H). This indicated a clear epithelium-to-fibroblast signalling.

In summary, in the presented model, CAFs primarily communicate with the overlaying epithelium via matrix-directed signals (Fig. 7A-B). In addition, cancer cells (especially E-basal-2 cluster) receive EGF, ncWNT and NRG signals from other epithelial cells, while MIF from epithelial cells and fibroblasts further supports cancer phenotype in HNSCC-like cluster 2. Suprabasal epithelial cells (cluster 4) are less engaged in the aforementioned signalling and instead present cell-cell junction interactions, while metabolic cluster 3 receives unique cancer– and barrier formation-related signals. In addition to self-sustaining FGF, PDGF and VEGF fibroblast signalling, cancer epithelium ensures the pro-tumour CAF identity is maintained by supporting CAF phenotype (TGFβ, NOTCH), survival and ECM production (MK/MDK), and establishing migration cues (ADGRL, SLITRK) (Fig. 7A).

**Figure 7.**
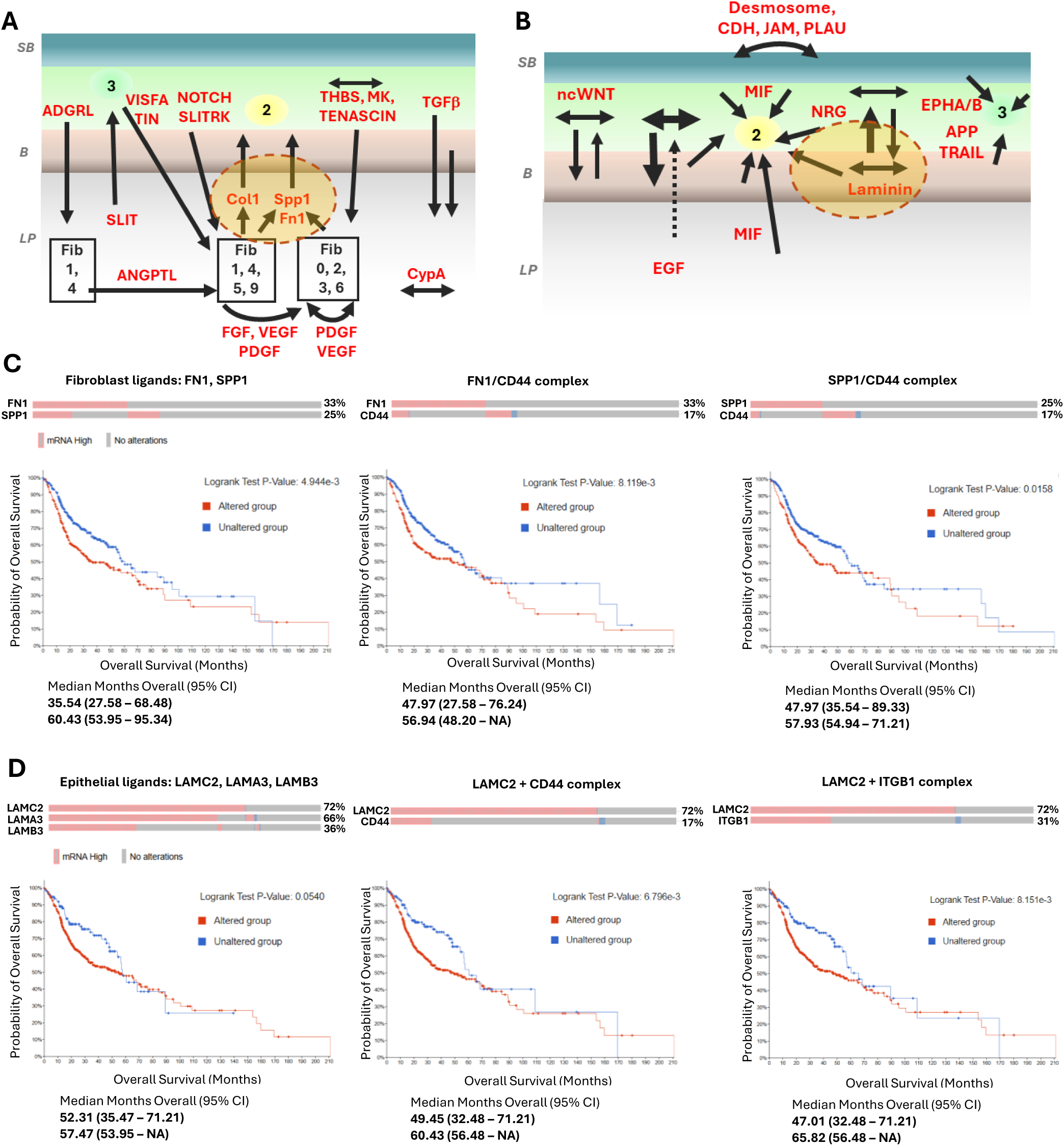
Clinical relevance of the cross-tissue signalling. (**A-B**) Graphical summaries of the signalling identified within the organotypic model. (**A**) Signalling between fibroblasts and epithelium and within the stromal cells. Specific CAF populations are indicated when relevant. (**B**) Main intra-epithelial signalling pathways within and between the basal and supra-basal layers with contribution from CAFs via paracrine MIF and EGF signalling. E-Basal-2 cluster is shown as the main receiver of tumour-sustaining signalling (cluster 2). Cluster 3-specific signalling is also indicated. SB, suprabasal layers; B, basal layers; LP, lamina propria. **(C-D)** Overall survival (OS) curves for 523 HNSCC patients with data available in TCGA PanCancer cohort with or without upregulation of genes (z-score ≥2.0, relative to normal samples). (**C**) The curves are shown for genes encoding fibroblast-derived ligands (FN1 and SPP1, left) and their complexes with CD44 epithelial receptor: FN1/CD44 (middle), SPP1/CD44 (right). (**D**) OS curves for genes encoding epithelium-derived ligands (laminins) and the primary identified ligand-receptor pairs: LAMC2/CD44 (right) and LAMC2/ITGB1 (right). Kaplan Meier Estimates were plotted using cBioPortal. P-values are calculated using log-rank test. The graph summary shows the difference in median months of OS between altered and unaltered groups. The frequency of each gene overexpression in the HNSCC cohort is shown above the graphs.

The signal from the matrix proteins (laminins, collagens, fibronectin and osteopontin) is the top signalling hub in the system, influencing the whole epithelium, but particularly HNSCC-like cluster 2. The importance of this signalling is highlighted by its activity having a clear impact on patients’ survival, as observed in the 523 samples from The Cancer Genome Atlas (TCGA) PanCancer panel with RNA expression data available. The upregulated expression of the top matrix-derived ligands (*FN1*, *SPP1*, *LAMB3*, *LAMC2*, *LAMA3*) and receptors (*CD44*, *ITGA2B1*) identified in the analysis is very common in the HNSCC tumours (up to 72% for *LAMC2*) and significantly decreases overall survival (Fig. 7C-D). This impact is in striking contrast to the lack of effect of the alterations in the identified autocrine signalling (EGF, ncWNT and MIF) (Suppl. Fig. 10). Therefore, interactions between epithelial cancer cells and stroma emerge here as a primary therapeutic target in HNSCC, and the characterised model is well equipped to test this further.

## 4. Discussion

Tissue-engineered models, including those of oral mucosa and epithelial cancers, present an animal-free, physiologically relevant and well-controlled platforms to study mechanisms of carcinogenesis and responses to therapies. However, their wider uptake is dependent on the increased confidence in how well they reproduce the in vivo tissues and how suitable they are for translational applications; answers to these questions can be addressed by extensive ‘omics profiling and comprehensive bioinformatics analyses (Gendoo, 2020). Here, the single cell reconstruction of the in vitro oral cancer model shows a well-developed, although not terminally differentiated, cancer epithelium, supported by stroma with highly heterogeneous fibroblast populations and extensive cross talk developed between the two tissues. To our knowledge, this is the first comprehensive characterisation of such a cancer model in single-cell resolution. However, there are a few examples where scRNA-seq has been applied to patient-derived (Um et al., 2024) and mouse model-derived organoids (Johansson & Ueno, 2021), and these data sets can validate the functionality of the fully in vitro grown organotypic cultures. In addition, the model characteristics can also be compared to data obtained from non-cancer organotypic and organoid cultures, including epithelial rafts exposed to human papillomavirus 16 (HPV16) (Bedard et al., 2023) and organoids derived from mouse tongue (S.-Y. Kim et al., 2025). More comprehensive literature addresses heterogeneity of in vivo tissues, both periodontium/oral mucosa (Caetano et al., 2021) and cancer (Arora et al., 2023; S. Kim et al., 2025; Kürten et al., 2021; Li et al., 2024; S. V. Puram et al., 2017), including the stepwise progression of HNSCC (Choi et al., 2023).

Considering the model was developed using established cell lines (HNSCC VU40T cells and NIH 3T3 mouse fibroblasts), the recreated oral cancer mucosa features highly comparable to the single cell landscape of the above-mentioned organoids and HNSCC tumours. As induced by air-liquid interface, VU40T cells developed stratified epithelium remarkably resembling the in vivo tissue of cancer origin with amplifying cells, heterogenous basal populations, supra-basal layer and TJ-enriched metabolism-oriented cells. These populations could be mapped due to the expression of keratins and junctional proteins and were localised within the model using immunostaining. The structure aligns with previous reports, which show unstructured, hyperproliferative tissues with reduced *KRT13* and clustered laminin-V expression, similar to in vivo HNSCC tissue (Gronbach et al., 2020). Although proliferating cells account for a small percentage of all keratinocytes after 20 days of culture, they are dispersed across whole tissue rather than restricted to para-basal layer.

The origins of all epithelial cells can be traced back to the amplifying cells, as expected in maturing epithelium. The basal populations differentiate via a precursor state and produce cells with basement membrane connections (E-Basal-1), while some of them undergo keratin to actin switch with activation of Rho-GTPases pathway (E-Basal-3 and E-LeadEdge-like). In many aspects, these two clusters show features of p-EMT/collective migration guided along collagen fibres, as described before for HNSCC tumours (S. V. Puram et al., 2017): decreased epithelial core markers, activation of Rho-ROCK-myosin pathway, coupled to ECM contact via DDR-1 (collagen-activated RTK). Unlike single-cell EMT, the cells in cluster 6 appear to maintain low E-cadherin-based adherens junctions (*VCL*, *CTNNB1*, *CTNNA1*), together with remodelled TJs evidenced by high *PATJ* expression, features typical in motile but cohesive tumour cell clusters (Parri & Chiarugi, 2010; S. V. Puram et al., 2017). In addition, the cluster is uniquely marked by high expression of KLF7, which is known to drive EMT, adhesion and migration programs across cancers (Cai et al., 2024). Therefore, E-LeadEdge-like cluster 6 could be the source of cells undergoing partial EMT and collective migration observed at the invasive front of carcinomas. However, clusters 5 and 6 are highly heterogeneous with only a small fraction of cells showing strong expression of the mentioned pEMT markers as well as vimentin (E-LeadEdge-like) or N-cadherin (*CDH2*) and *SNAI2* (E-Basal-3) therefore preventing any definite conclusions regarding their functions. There is also a possibility that a partial EMT transcriptional programme could be induced by organotypic culture conditions as previously reported in skin cultures although this was observed when using non-transformed cells (Stabell et al., 2023).

By comparison, E-Basal-2, although strongly associated with ECM pathways and COL17A1, shows lower enrichment for laminins than E-Basal-1, indicating more para-basal location. This cluster is the main hub for HNSCC-overexpressed TFs (*NFKB1*, *MYC*, *E2F*, *ETS*) and HNSCC signalling pathways as later confirmed by its engagement in the crosstalk with other epithelial clusters and with cancer stroma. However, some of the cells escape cancer-like programme and differentiate into keratin-, desmosome-, and TJ-rich supra-basal layer (E-Suprabasal).

After ranking the clusters according to the maturation trajectory, the gradient of TF expression can be easily recognised. The first maturation change marks exit from the proliferating state by switching on *SOX4* expression. SOX4 activity is traditionally linked to proliferation, however, in ora cancer it has been implicated in suprabasal differentiation and barrier formation (Moreno, 2019). In the in vitro HNSCC model this could be reflected by E-Cycling cluster reliance on classical transcription factors for proliferation (MYC, E2Fs), while SOX4 operates by guiding differentiation, remodelling plasticity and metabolic adaptation in post-mitotic cells. Next, the change from less differentiated to most differentiated (E-Suprabasal, E-Basal-1) populations is associated with gradual upregulation of *EHF* and *PITX1* together with decrease in *RUNX1* expression in general and change from MYC to MXD1 as a partner for MAX transcription factor supporting the differentiation of supra-basal epithelium, in particular. It correlates with a switched-on expression of *HES2* and *PRDM1* in the E-Suprabasal cluster. Majority of these transcription factors are well known in epithelial biology and craniofacial development. Similarly, AP-1 and KLF transcription factors are major regulators of epithelium development with expression changes frequently linked to cancer progression. Specifically, KLF4 down-regulation is linked to impaired differentiation while KLF5 expression is needed for sustaining basal-like activity in oral cancer (McConnell et al., 2007). While AP-1 factors are key for epithelium maturation, wound healing and inflammatory responses, their activation is controlled and transient in physiological conditions (Eckert et al., 2013). In HNSCC, AP-1 activation is constitutive and persistent, driving cancer development and EMT-like states (Wang et al., 2021). These cancer-like transcriptional regulations are recapitulated in the in vitro model presented here. Imposed on it is the basal cell programme, centred around TP63 TF, which in the in vitro model is supported by glucocorticoid receptor (*NR3C1*) and *CEBPB* expression.

In parallel to the canonical suprabasal differentiation, an alternative epithelial maturation trajectory was identified in cluster E-Metab, emerging directly from proliferating cells. This pathway is characterised by activation of a tight-junction–enriched, metabolically active epithelial programme with non-classical keratin expression (KRT7/8/81). This may suggest a barrier-focused, energetically demanding epithelial state governed by unique set of transcription factors either specific (NFIA, MEIS2, CLOCK), or shared with E-Cycling (NR6A1, GATA6, NFIC) or E-suprabasal (ELF3, GRHL2) clusters. This could be representative of cancer tissue plasticity or an advantage of scRNA-seq data to facilitate identification of alternative trajectories (S. V. Puram et al., 2017). However, such population has not been previously described in the single cell-based studies therefore the observed state could also be enhanced by in vitro culture at air-liquid interface. It has been previously shown that epithelial barrier integrity and TJ organisation increase in organotypic cultures compared to native tissues (Stolte et al., 2020).

Tumour microenvironment is a well-recognised contributor to carcinogenesis, it can modify tumour progression and metastasis, cancer cells responses to therapies/resistance to therapies, as well as support for immune tissue component and vascularization (Chhabra & Weeraratna, 2023). CAFs are the main functional cells in the tumour stroma but their contributions and functions are still poorly understood due to their remarkable heterogeneity, which scRNA-seq methodology has helped to recognize in the recent years (Bensa et al., 2023; Cords et al., 2023; Liu et al., 2025; Sahai et al., 2020). This heterogeneity results both from their varied origins and the signaling received from tissue matrix and other cells (Chhabra & Weeraratna, 2023). In our model, the origin of fibroblasts is single and well defined (mouse embryonic NIH3T3 cells). Yet the in vitro tissue culture provided enough stimulus to allow for their differentiation into all CAFs’ subtypes described so far: mCAFs, iCAFs (including ifnCAFs), dCAFs and tCAFs. The exact subpopulation markers are different in some cases, most likely due to the mouse origins, but the functionalities are well reproduced. For example, the tissue lacks the main myCAFS (myofibroblastic) markers, a-SMA or POSTN, however, clear matrix-producing populations are identified.

The study demonstrates for the first time, the differentiation from a single fibroblast starting population into multiple CAF subtypes within an organotypic model and with lineage evidence. Previous studies either directly used patient-derived CAFs (Strating et al., 2023) or reported general activation of normal fibroblasts towards CAFs (Kim et al., 2024). What appears to be key for achieving heterogeneity in vitro is the differentiation pathways which originate from the dividing cells. Only populations newly differentiated within the matrix were able to develop into one of the described CAFs’ components. The second source identified by the pseudotime analysis resulted in fibroblast populations with mostly metabolic characteristics. These are most likely the fibroblasts that were originally seeded within the collagen and underwent terminal differentiation. However, it is not clear whether they also contribute to the tCAFs population.

Similarly, only the newly differentiated CAF populations engage in intra– and inter-tissue crosstalk while the originally seeded fibroblasts remain quiet. Most importantly, this study demonstrates that physiological communication between cancer cells and stroma can be reproduced remarkably well in organotypic cultures. As expected, fibroblast/CAFs predominantly depend on FGF and PDGF (in addition to ANGPTL and CypA) signalling to form self-sustaining loops within the stroma, with TGF-β ligands coming exclusively from epithelium despite epithelium lacking the receptors. This is supported by Notch, ADGRL, THBS, SLITRK, MDK and Tenascin epithelial signals. The autocrine/paracrine signals received by epithelium involve non-canonical WNT, EGFR, NRG and MIF pathways. Apart from MIF, most of this signalling is produced within epithelium with only small contribution from fibroblast. However, the most prominent stimulus received by cancer cells, especially by E-Basal-2, HNSCC-like cluster, is matrix-generated, either by stroma (collagens, fibronectin and osteopontin/SPP1) or epithelium (laminins). All components of this signalling are overexpressed in squamous oral tumours and the crosstalk between collagens and CD44 or integrins has been recognized before in scRNA-seq analysis of human tumours (Choi et al., 2023; S. V. Puram et al., 2017). Although FN1 and SPP1 fibroblast ligands were not mentioned in these studies, we show here their alterations have significant impact on overall patient survival in HNSCC PanCancer Atlas cohort and clearly surpassing the effects of the components of the paracrine pathways. This emphasizes the relevance of structural stromal proteins in maintaining the bulk of epithelial tumour and presents them as potential therapeutic targets.

Targeting tumour-stroma interactions has been pursued for some time although none of the solutions have been clinically implemented (Chitty & Cox, 2025). Our study provides evidence that organotypic models are suitable platforms for pre-clinical validation of such therapeutic approaches. Both fibronectin and osteopontin are already of interest for cancer therapy, however most applications propose them as delivery tags for other therapeutics (i.e. cytokines, radionuclides) to reach and concentrate in tumour bed (Kähler et al., 2025), or modulators of immunotherapies (Di Nitto et al., 2024). This study supports the newer ‘stromal normalization’ approach. Reducing fibronectin assembly (using PEG-FUD) has been shown to suppress tumour growth and progression in breast cancer mouse model (Gari et al., 2025) while volociximab/M200 antibodies can be used clinically to target main fibronectin receptor, α5β1 integrin (Mateo et al., 2014). In addition, the specificity of the targeting can be increased by the presence of tumour-associated fibronectin isoforms (EDA and EDB) (Kumra & Reinhardt, 2016). Direct SPP1 therapeutics are less developed although experimental neutralising antibodies exist (Bandopadhyay et al., 2014). A recent HNSCC study shows that targeted suppression of SPP1 inhibits invasion/metastasis and cisplatin resistance in NRF2-hyperactivated, cisplatin-resistant HNSCC (Kawabe et al., 2025). This context is particularly relevant for our model where majority of tumour cell populations exhibit high NRF2 overexpression. In addition, we previously showed that targeting the oxidative response pathway in several HNSCC cell lines (including VU40T) is beneficial, at least in in vitro studies (Gleneadie et al., 2021).

Oxidative stress response pathway emerges in the model as major characteristic, known to play key role in resistance to cancer therapies. This characteristic is primary associated with E-Prec and E-Metab clusters, while several markers linked to chemoresistance in HNSCC tumours are enriched in E-Cycling population.

Therefore, the model presents with multiple, in vivo-like characteristics suitable for cancer modelling and drug testing, including cell population enriched in HNSCC markers and pathways (E-Basal-2), populations with prominent therapy resistance pathways, cancer-stroma interactions of translational potential and cells possibly forming an invasive front. Including immune cells in the model would broaden the scope for exploring and harnessing these interactions.

## 5. Conclusions

This study demonstrates the high value of a non-animal, human-based platform for functional and therapeutic testing, especially for interrogating tumour-stroma signalling in a physiologically relevant tissue context. In addition, it substantially advances our understanding of epithelial biology by uncovering subtle cellular states and reconstructing transcription factor-driven maturation trajectories associated with differentiation, barrier formation and metabolic adaptation in cancer epithelium. Finally, the study highlights cancer-stroma interactions as clinically relevant therapeutic targets and establishes a robust experimental system for the evaluation of existing and future treatment strategies.

## 6. Experimental Section

### Cell culture and generation of organotypic cultures using BECDs

Air–liquid interface (ALI) cultures were established using buoyant epithelial culture devices (BECDs) fabricated via fused filament fabrication (FFF) using polypropylene, following the protocol described by Hewitt et al., 2022. Murine 3T3 fibroblasts (ECACC 85022108) and human female VU40T tongue-derived oral squamous cell carcinoma cells (Prof. H. Joenje, VU University Medical Center, Amsterdam) were cultured in complete Dulbecco’s Modified Eagle Medium (DMEM; Gibco, UK) supplemented with 10% foetal bovine serum, 100 U/ml penicillin, 100 μg/ml streptomycin (Gibco, UK), and 1 mM L-glutamine (Thermo Fisher, UK). The cells were regularly tested for Mycoplasma and used at passage 18 and 22. Hydrogel preparation was carried following the previously published protocol (Hewitt et al., 2022). Briefly, 1 ml of 3 mg/ml rat tail collagen I (Thermo Fisher) was mixed with 100 μl of 10x DMEM (Gibco) and 100 μl of 10x reconstitution buffer (prepared by dissolving 1.1 g NaHCO₃ and 2.385 g HEPES in 50 ml dH₂O; both from Sigma-Aldrich, UK). The pH was adjusted by adding 1 M NaOH dropwise until a bright-pink solution was achieved. 3T3 fibroblasts were added to the hydrogel at a final concentration of 5 x 10⁴ cells/ml. A total of 1.2 ml of the hydrogel-cell suspension was dispensed into each BECD positioned in a 6-well plate (NUNC, UK). The hydrogels were incubated at 37 °C, 5% CO₂ for 90 minutes to allow gelation, followed by the addition of 3 ml of complete DMEM per well. After 24 hours, 8 x 10⁵ VU40T cells were seeded on top of each hydrogel. Cells were allowed to attach for 30 minutes before adding an additional 3 ml of culture medium to each well. Submerging rings were then placed over the BECDs and the cultures were incubated for 72 hours to allow epithelial monolayer formation. After the initial submersion phase, the rings were removed, allowing the BECDs to float and establish an air–liquid interface with the keratinocyte layer exposed to air. Cultures were maintained for 20 days, with media replaced every other day.

### Single cell RNA sequencing

Tissue dissociation: to isolate single cells from the 3D hydrogel cultures, enzymatic and mechanical dissociation of the gels was performed. Two hydrogel constructs per experiment were processed in GentleMACS™ C Tube (Miltenyi Biotec, Germany). An enzyme digestion mix consisted of 600 µl of dispase II (10 mg/ml, Sigma), 1333 µl of collagenase I (4 mg/ml) (Gibco, US), 20 µl of DNase I (1 mg/ml, Merck), and 46.6 µl of DMEM. The hydrogel was immersed in this mix and processed for 21 minutes at 37 °C using the GentleMACS™. The resulting cell suspension was passed through a 40 µm cell strainer to remove large aggregates. The filter was then washed with 1 ml of pre-warmed 0.25% (w/v) trypsin in 1 mM EDTA and incubated at 37 °C for 5 minutes. The suspension was mixed by pipetting and incubated for an additional 5 minutes. Visual inspection was performed to assess residual debris. The suspension was combined with the earlier filtrate, and cell concentration was estimated by an automated counter (Luna-FX7, Logos Biosystems, South Korea). Cell debris was removed from the suspension following the manufacturer’s protocol (Debris Removal Solution, Miltenyi Biotec, Germany). The final cell pellet was resuspended in 1 ml of chilled 0.04% BSA in PBS and processed immediately.

The GEM (Gel Bead-in-emulsion) and single cell library was prepared using 10X Genomics Chromium Single Cell 3’ Library & Gel Bead Kit v3.1. The library was sequenced using NovaSeq 6000 (100 cycles) and SP Reagent Kit v1.5 in the Birmingham Genomics facility.

### Data Processing & Genome Alignment

The FastQ files produced from each sample were run through the nf-core/scrnaseq v. 4.0.0 pipeline using the 10x Cell Ranger aligner. Due to the presence of both human and mouse cells in the model setup the reads were aligned to both genomes using hg38 and GRCm39 for respective analysis of epithelial cells and fibroblasts. For the hg38 genome, a Cell Ranger reference was constructed using the 10x platform as described previously (Zheng et al., 2017). For the GRCm39 genome, the CellRanger reference was downloaded directly from the 10X Genomics Cell Ranger v4.0.0 using 10X Genomics Cloud Analysis. 10X Cell Bender outputs, which remove intrasample ambient RNA contamination, converted to Seurat in RDS format from the pipeline were loaded into Seurat (v.5.3.0) in R v.4.4.1. To quality control each sample, we excluded cells containing > 20% mitochondrial reads, fewer than 300 transcripts, and detected gene counts (nFeatures) which differ by 5 median absolute deviations (MAD) from the log-transformed distribution of cellular gene counts (nFeatures) (Germain et al., 2020; Luecken & Theis, 2019). DoubletFinder (v.2.0.6) was then run on each sample separately to remove doublets in an automated fashion (McGinnis et al., 2019). The optimum number of principle components (PCs) used to construct the uniform manifold approximation and projection plots (UMAP) and for clustering of the integrated datasets for each separate alignment was determined by JackStraw score analysis performed on the top 20 PCs to select the optimum number of PCs. The optimum resolution for clustering was determined by the construction of a clustree (v. 0.5.1) (Zappia & Oshlack, 2018) for each alignment at resolutions between 0.1 and 1.5 at intervals of 0.1. The most stable level of which in addition to cluster markers identified were used to inform the resolution used for clustering (0.3 for human only, 0.6 for mouse only data).

### Multi-species alignment and cell annotation

Data from both genome alignments were combined and each cell was classed as ‘human’ or ‘mouse’ by aggregating transcript counts and assigning cell barcodes based on a ≥ 70% identity cutoff from each respective alignment. A small number of cell barcodes that did not meet this threshold were assigned as ‘mixed’ and removed from subsequent analysis. The resulting 9520 human cells and 22293 mouse cells were re-clustered as described above and assessed by the presence of manually curated lists of markers widely accepted for the two lineages. One of the mouse clusters (cluster 13) did not align with fibroblast marker assigned averages and the cells were also removed from the subsequent analysis, resulting in 21,737 fibroblast cells.

Heatmaps, dotplots and violin plots were constructed from predefined marker lists, using the Seurat objects derived from both species separately using R package ComplexHeatmap (v.2.22.0) and the Dotplot and Vlnplot functions from Seurat, respectively. The pre-defined marker lists are indicated in each figure. Feature plots for markers of interest were also generated using the FeaturePlot function from Seurat.

### Over-representation analysis (ORA)

The top 200 (or maximum number if below 200) positively expressed markers from each cluster were derived from running the Seurat FindAllMarkers function on the counts for each individual integrated fibroblast and epithelial Seurat object (logFC > 0.25 and expressed in minimum 25% cells per cluster, p_val_adj < 0.05). These were then run on DAVID (Knowledgebase v2025_1) (Sherman et al., 2022) using gene sets pertaining to REACTOME, Wikipathways or combined KEGG/REACTOME/Wikipathways as indicated in the results. The top 10 enriched pathways for each cluster with an FDR < 0.05 were combined and manually curated to avoid redundancies and plotted. The representative genes were selected from the most enriched and significant groups and manually curated to show gene expression patterns across the clusters.

### Trajectory Analysis

Monocle3 v 1.2.46 (Trapnell et al., 2014) was used to construct a pseudotime trajectory for both epithelial and fibroblast clusters separately using default parameters on their UMAP embeddings, respectively. A root cluster of 7 was chosen for the epithelial clusters after UMAP overlay inspection of mitogenic factors. Root clusters of 0 and 9 were chosen as the root clusters to construct a pseudotime trajectory for the fibroblast clusters.

### Cell-cell communication analysis

Human homologues (Ensembl v. 115) were mapped onto the raw expression data of the separate mouse fibroblast clusters using bioMart, merged with the raw counts of human only epithelial clusters and then re-log-normalised using the Seurat NormalizeData function. Cell communication analysis on the log-normalised counts of this merged object was performed using R package CellChat (Jin et al., 2025) with default parameters. The CellChatDB human was used for analysis. Outgoing and incoming cluster and signalling pathway patterns were analysed using the R package NMF v 0.28 and visualised with ggalluvial. The NMF Cophenetic and silhouette scores obtained through the CellChat selectK function were used to infer the number of patterns for all clusters in both signalling directions.

### TCGA data

Gene expression and patient survival data were retrieved from The PanCancer Genome Atlas (TCGA) Head and Neck cohort (n=523) via cBioPortal for Cancer Genomics (Gao et al., 2013) and included the percentage of samples with each gene alteration (mRNA Expression z-score threshold ± 2.0 relative to normal samples), as well as Logrank test p-values for overall survival differences between samples with and without alterations. Kaplan-Meier Estimates were plotted using cBioPortal.

### Histology

Hydrogels were fixed in 10% neutral buffered formalin overnight at room temperature, then washed three times with phosphate-buffered saline (PBS). Fixed hydrogels were dehydrated through a graded ethanol series (10–100%, 10 min each), with two additional immersions in 100% ethanol. Samples were cleared in xylene (2 × 10 min) and infiltrated with paraffin wax at 60 °C overnight. The paraffin wax was replaced with fresh wax, and samples were incubated for a further overnight period. Sections of 4 µm thickness were cut using a rotary microtome (RM2035, Leica, Germany), mounted on Superfrost Plus™ glass slides (Sigma-Aldrich, UK), and incubated at 60°C for 1 h. Slides were dewaxed in two successive xylene washes and subsequently stained with either H&E or immunofluorescent dyes.

### H&E staining

Sections were stained with Gill’s Hematoxylin III (Sigma-Aldrich, UK) for 4 minutes to visualise nuclei, then rinsed under running tap water. To enhance nuclear contrast, slides were immersed in 0.3% acetic acid for 30 seconds, followed by differentiation in 0.3% hydrochloric acid in 70% ethanol for 30 seconds. Nuclear staining was enhanced in Scott’s tap water for 2 minutes, then rinsed in tap water. Eosin solution (Sigma, UK) was applied for 1 minute to stain the cytoplasm and extracellular matrix. Slides were dehydrated through graded ethanol solutions (10-70%, 95%, and 100%, 2 minutes each), cleared in xylene (2 × 5 minutes), and mounted with DPX mounting medium (Sigma, UK). Stained sections were imaged using a brightfield Primotech HD Transmission Microscope Monocular (Zeiss).

### Immunofluorescence

Additional organotypic cultures were grown as biological replicates and used for validation by immunofluorescence staining. Paraffin-embedded tissue sections were deparaffinised by two 5-min washes in xylene, followed by a 5-min wash in 1:1 (v/v) xyline:ethanol solution. The slides were then rehydrated through a graded ethanol series (100%, 95%, 80%, 70%, and 50%) for 3 min each and rinsed in dH2O. Subsequently, the sections were washed in PBS three times for 5 min each. Antigen retrieval was performed using sodium citrate buffer (pH 6.0; Sigma, S4641). The sections were heated in a microwave oven for 10 min at medium power and allowed to cool in dH2O for 30 min at room temperature. For permeabilisation, the sections were incubated in 0.2% Triton X-100 (Sigma, HFH10) in PBS for 5 min at room temperature, followed by three washes in PBS for 3 min each. Non-specific binding was blocked by incubating the sections in 3% bovine serum albumin in PBS for 1 h at room temperature. After blocking, the slides were incubated with primary antibodies overnight at 4 °C. Then, then slides were either mounted directly using a DAPI-containing mounting agent (Abcam, AB104139) or incubated with secondary antibodies (Donkey anti-rabbit Alexa Fluor 488 (Invitrogen) and anti-mouse Texas Red (Jackson Immunoresearch Laboratories, USA) at 1:600 dilution for 1.5 h at room temperature before mounting. The list of primary antibodies used and their dilutions are included in Suppl. Table 1.

---

**Supplementary Figure 1.**
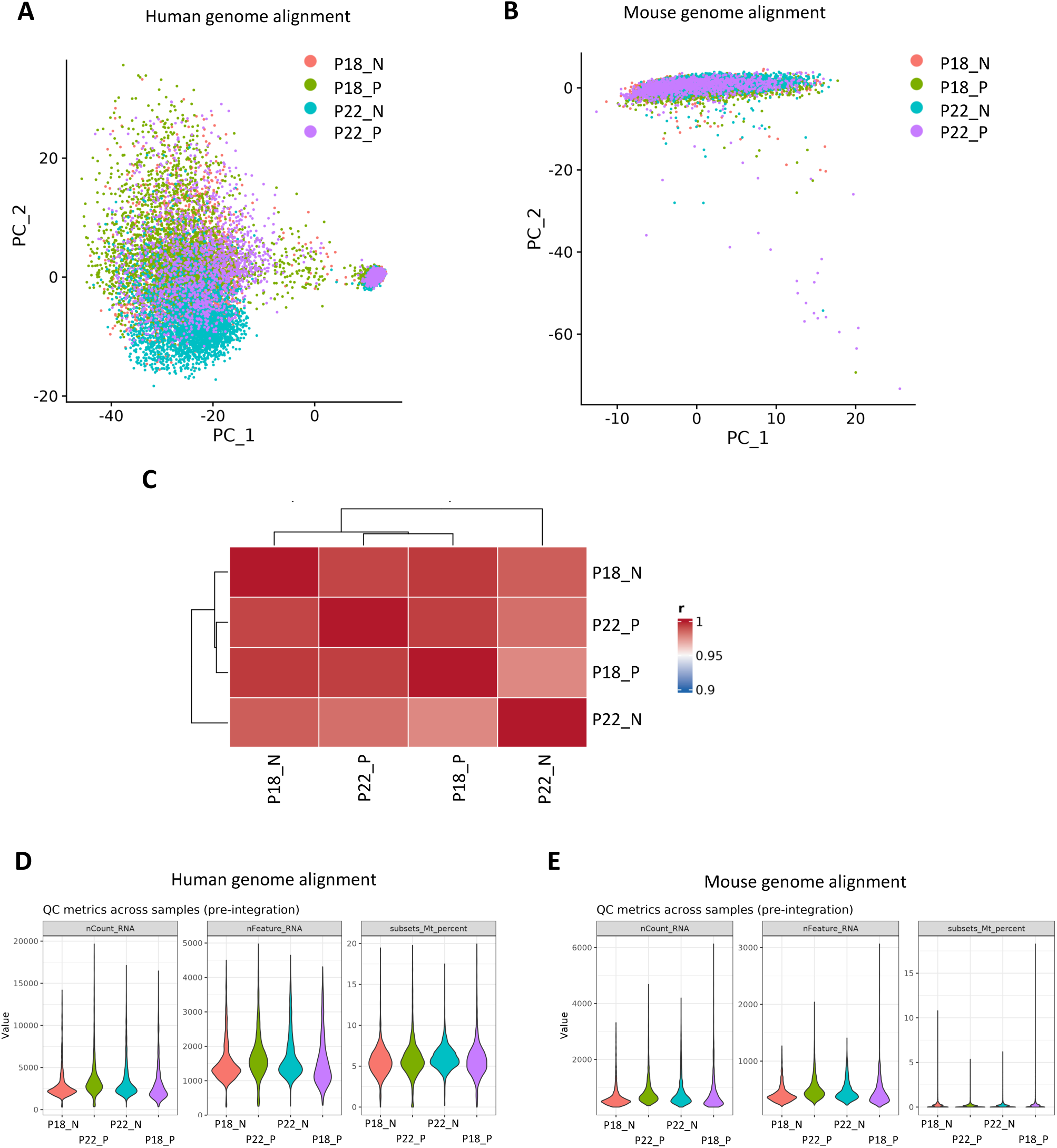
Assessment of inter-sample batch effects pre-integration. (**A-B**) PCAs of pseudobulked whole sample transcriptomes P18_N, P22_P, P18_P and P22_N prior to species filtering and integration on the (**A**) human genome alignment and (**B**) mouse genome alignment. **(C)** Hierarchically clustered heatmap showing pairwise similarity between samples’ pseudobulked whole transcriptomes P18_N, P22_P, P18_P and P22_N before integration. Colour intensity indicates similarity (range Pearson’s r ∼0.9–1.0), with darker red representing higher similarity. **(D-E)** QC metric violin plots pre-integration of all four samples showing per-sample distribution of number of transcripts (nCount_RNA), number of genes (nFeature_RNA) and percentage mitochondrial reads per cell detected (subsets_Mt_percent) for samples aligned to hg38 (D) and GRCm39 (**E**), pre-species filtering.

**Supplementary Figure 2.**
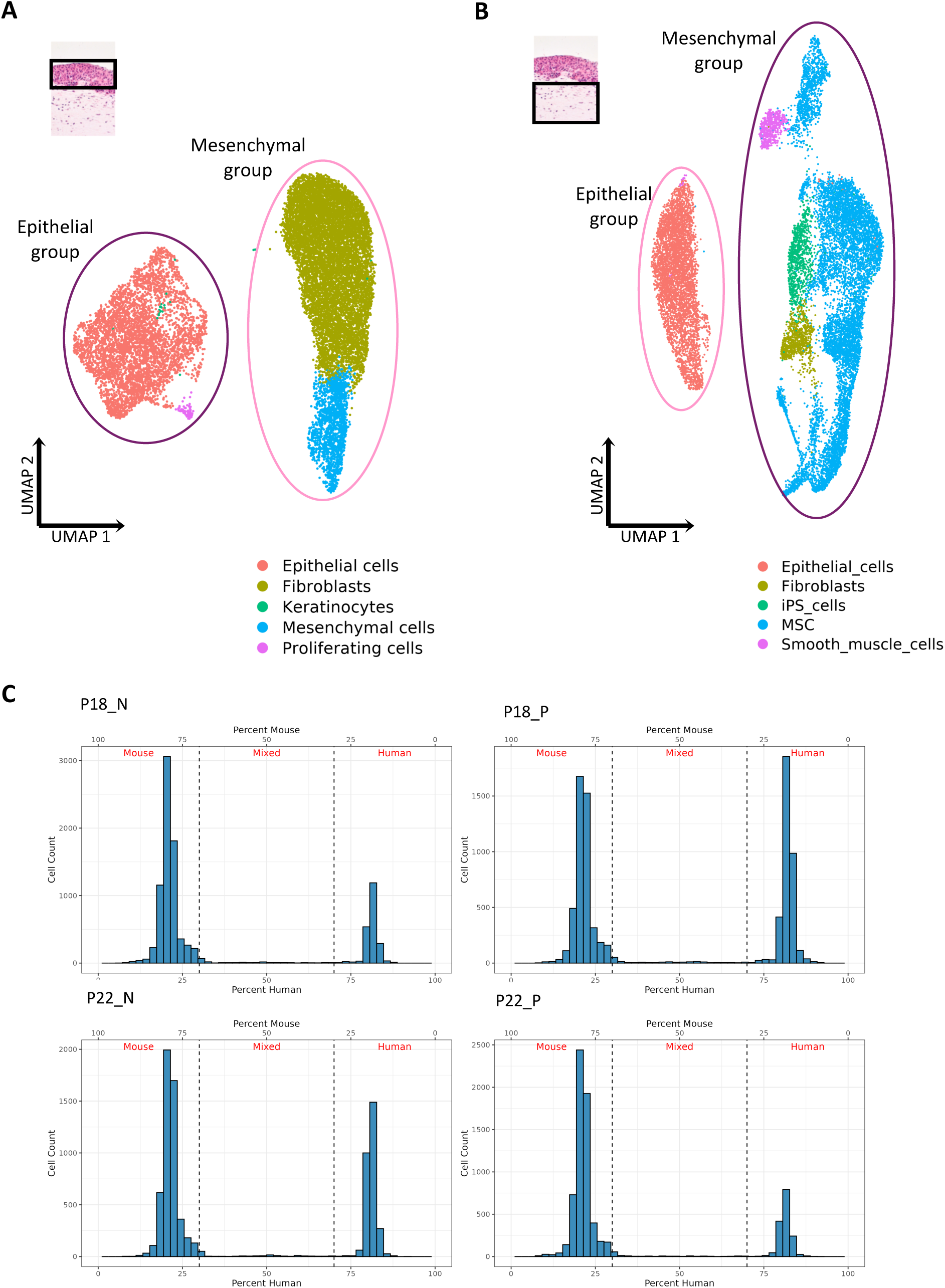
Initial clustering of integrated samples. (**A-B**) UMAPs of CCA-integrated samples without species-specific cell origin filtering by using the human alignment annotated with GPTCelltype (model: GPT-4o) (**A**) and using the mouse alignment annotated with SingleR (**B**). **(C)** Histograms showing the distribution of per-cell species assignment scores for samples P18_N, P18_P, P22_N, and P22_P, calculated after joint alignment to combined human and mouse reference genomes. The x-axis represents the percentage of human versus mouse reads per cell and the y-axis shows cell counts. Vertical dashed lines indicate ≥70% species-specific transcript thresholds distinguishing mouse, mixed and human cells.

**Supplementary Figure 3.**
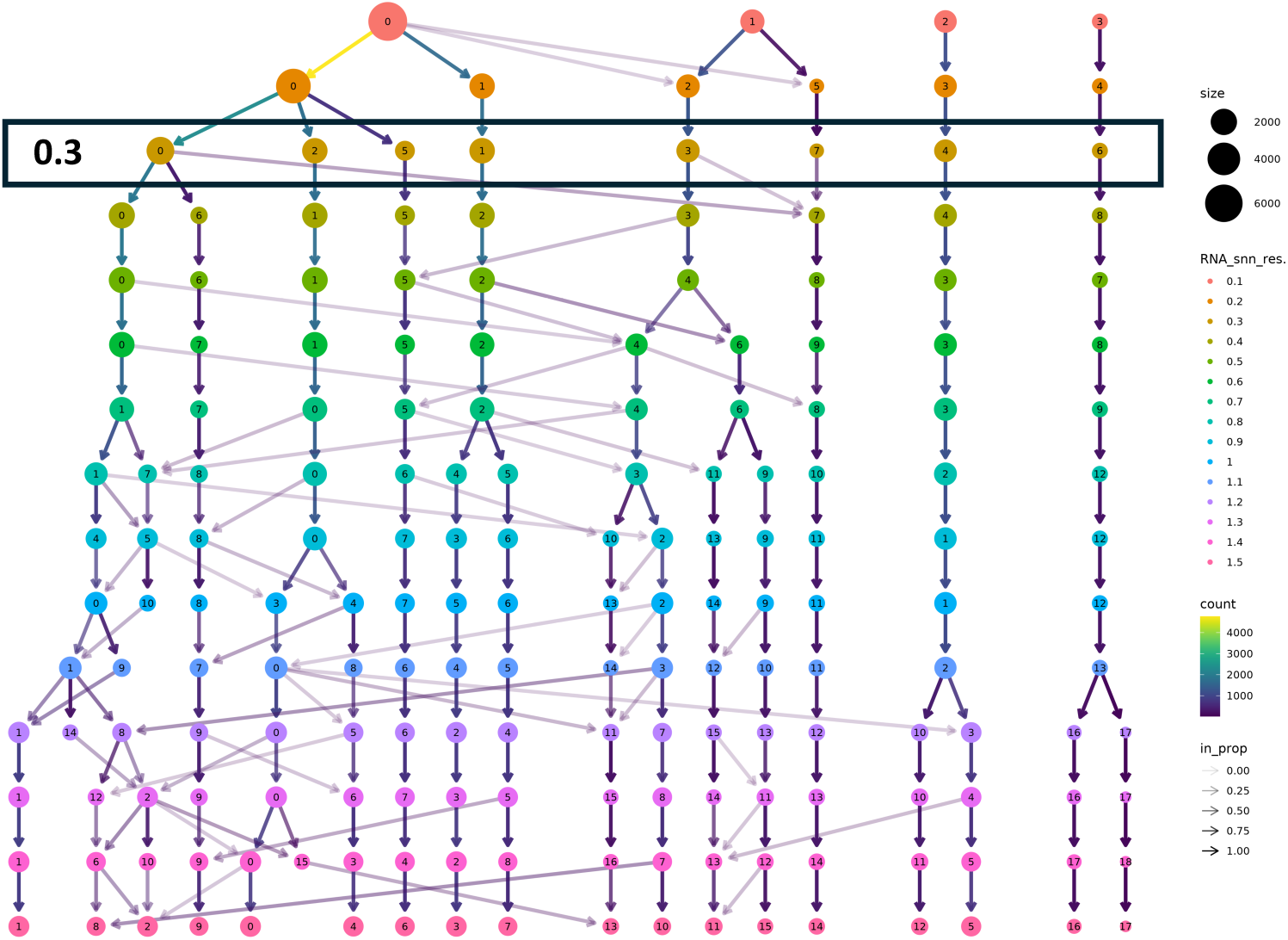
Assessment of cluster stability and purity using Clustree to determine the optimum cluster resolution for the integrated human alignment post-species-specific cell filtering and reclustering (yielding 9,520 human cells) using the top 10 PCs. 0.3 was determined as the most stable cluster resolution.

**Supplementary Figure 4.**
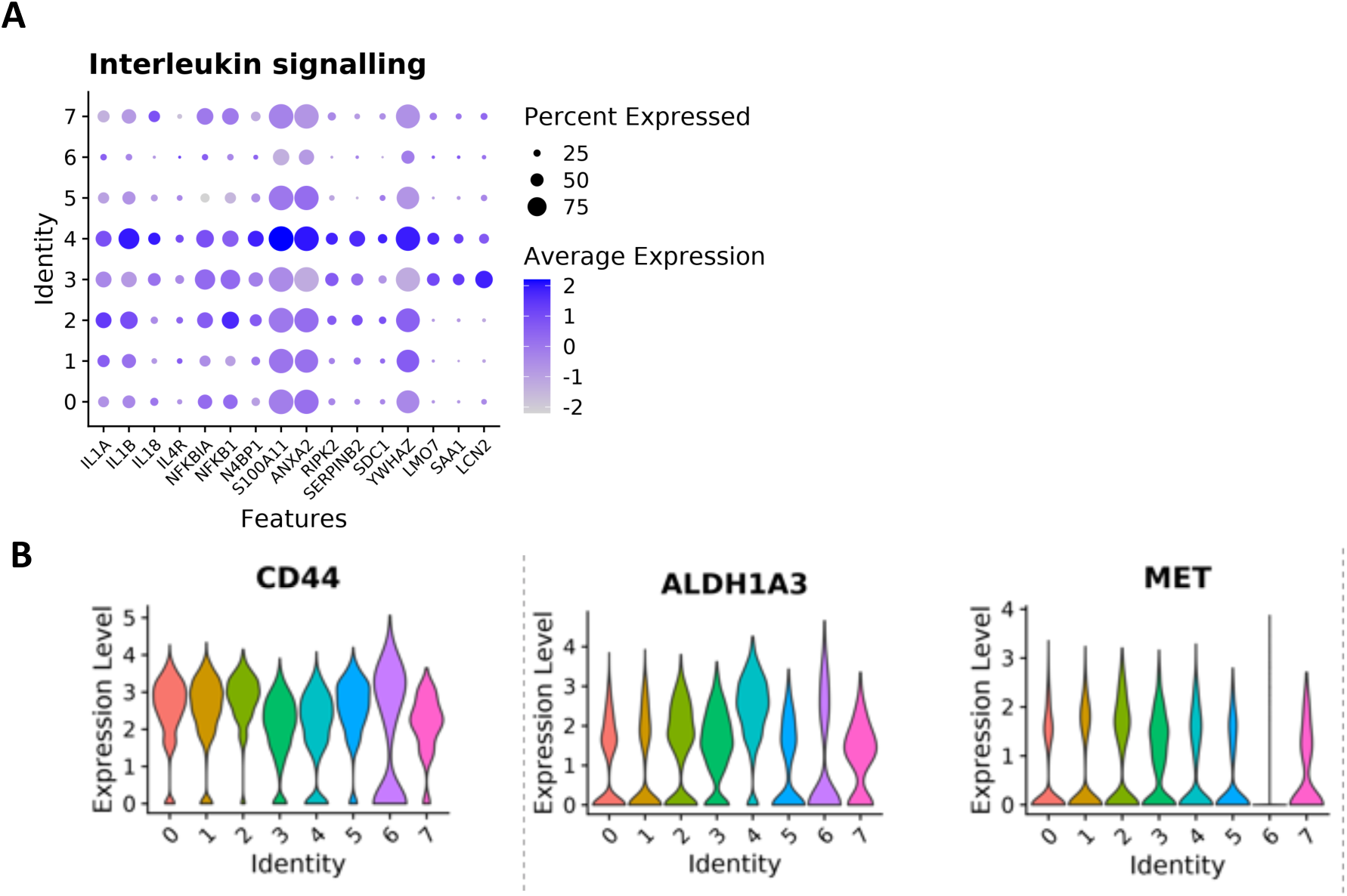
Additional analyses of the cancer component of the model. (**A**) Dot plot showing relative expression of genes encoding interleukins and involved in innate immune signalling across epithelial clusters. **(B)** Violin plots of relative gene expression for genes previously implicated as markers of cancer stem cells in HNSCC.

**Supplementary Figure 5.**
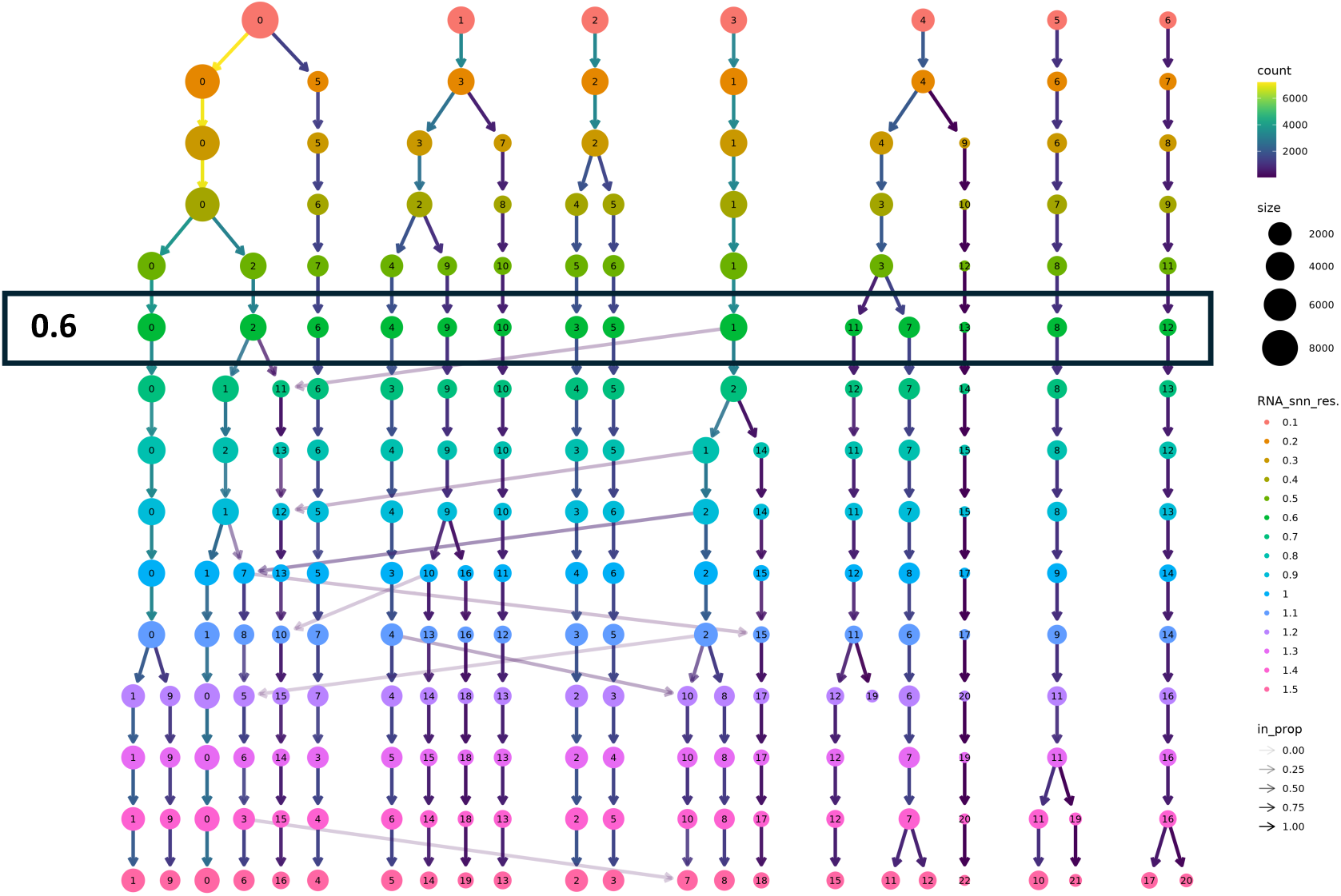
Assessment of cluster stability and purity using Clustree to determine the optimum cluster resolution for the integrated mouse alignment post-species-specific cell filtering and reclustering (yielding 22,293 mouse cells) using the top 20 PCs. 0.6 was determined as the most stable cluster resolution.

**Supplementary Figure 6.**
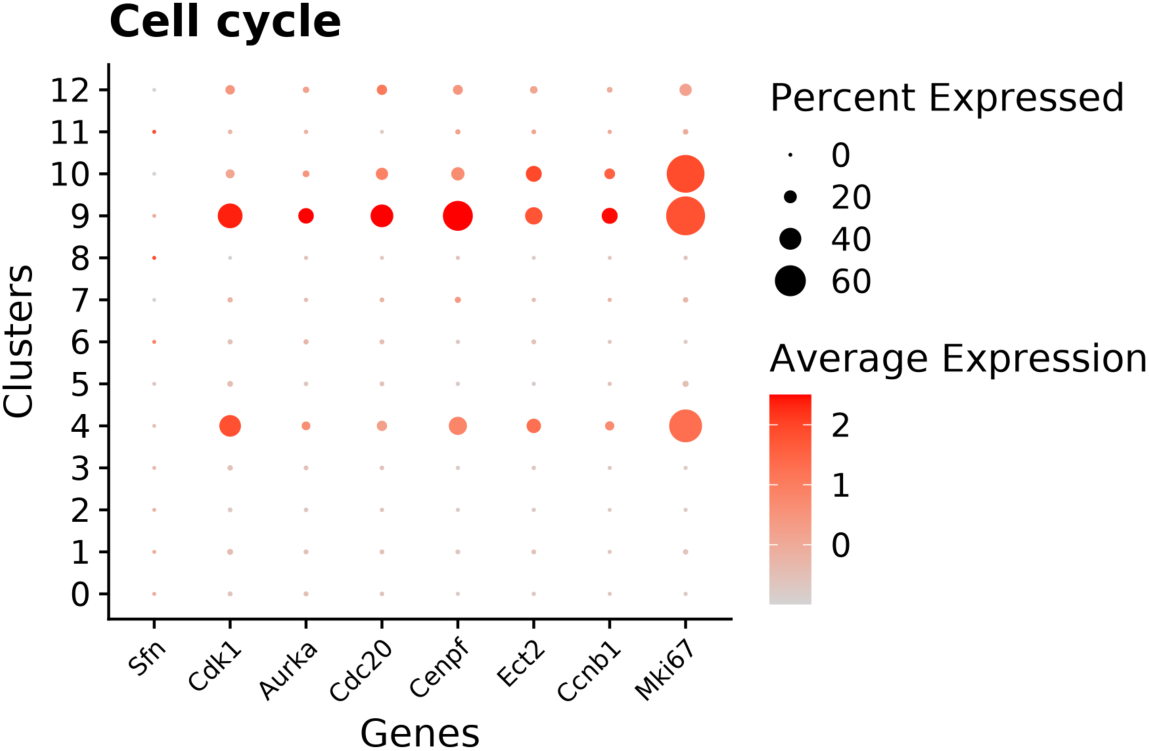
Gene expression dot plot for genes associated with cell cycle and proliferation in fibroblast clusters.

**Supplementary Figure 7.**
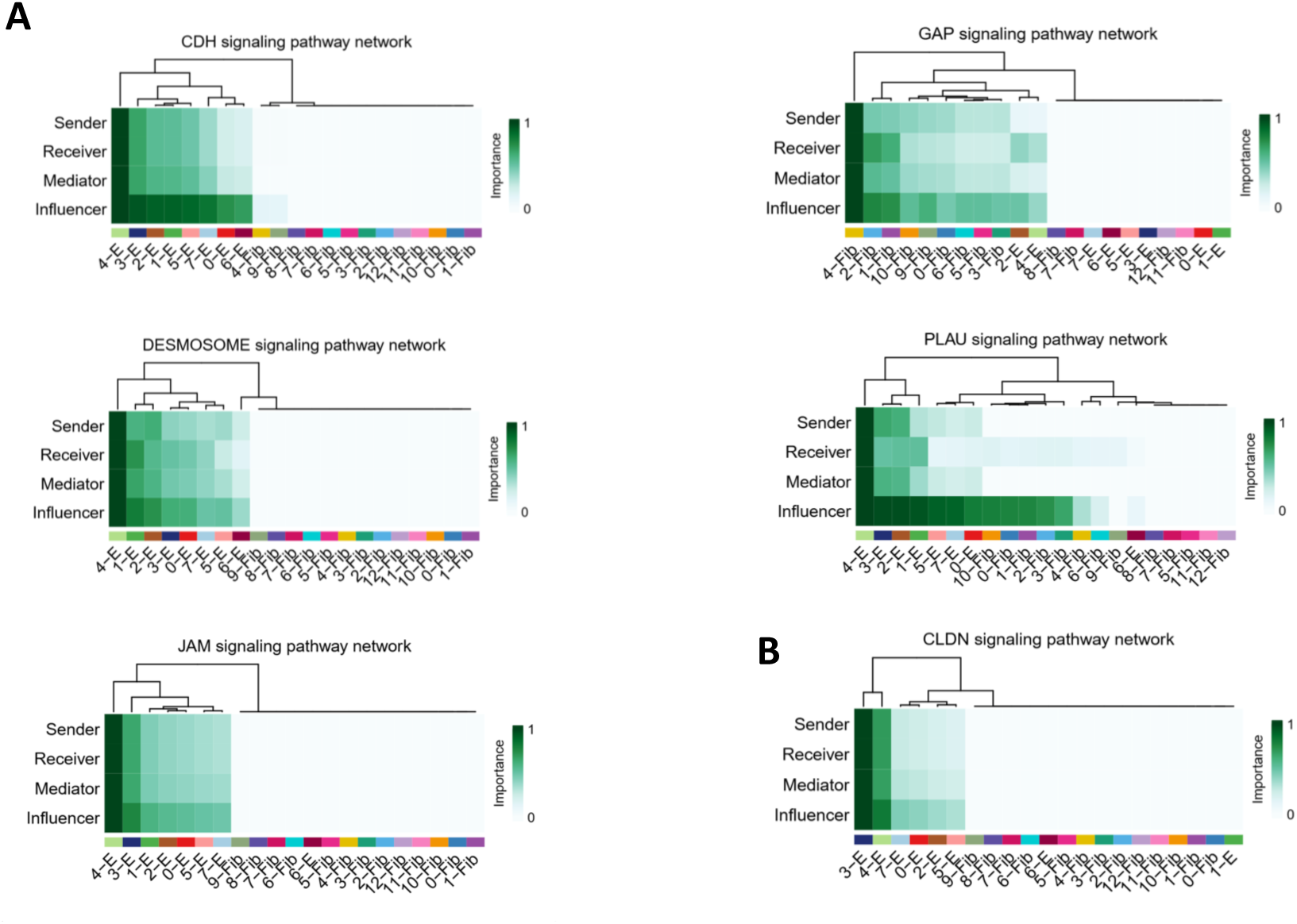
Intra-epithelial signalling characteristic for Pattern 3 of outgoing signalling (shown in Fig. 6B). (**A-B**) Relative contribution of each cell cluster in the intra-epithelial signalling involving direct cell-cell contacts within Suprabasal cluster 4 (**A**) and Metabolic cluster 3 (**B**).

**Supplementary Figure 8.**
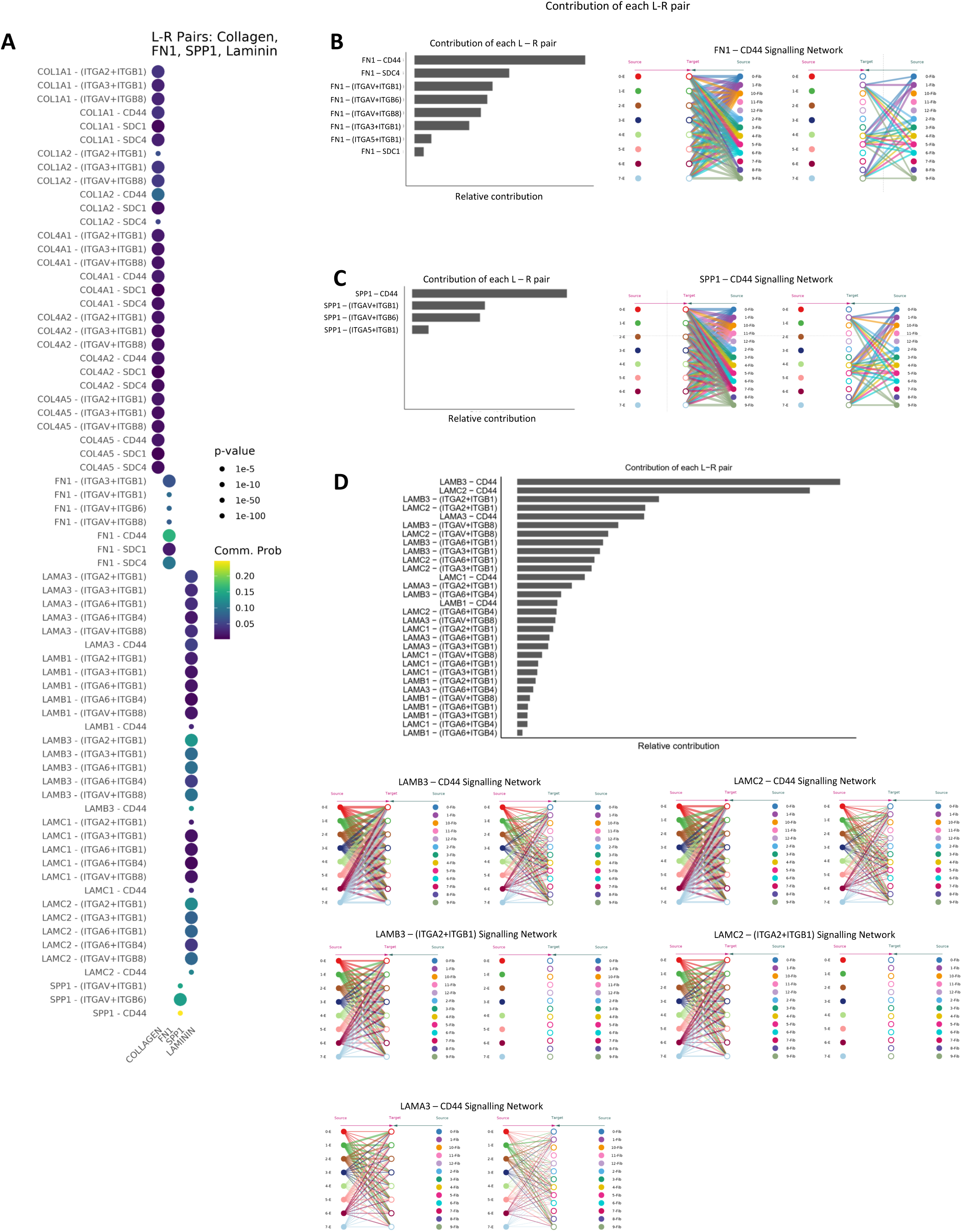
Detailed characterisation of signalling between matrix-derived ligands (Collagens, FN1, SPP1 and Laminins) and their associated receptors. (**A**) Dot plot showing relative probability of the listed ligand-receptor pairs. **(B-D)** Relative contribution of the main receptor interactions (bar graph) and the target-source signalling network for the most common interaction with FN1 (B), SPP1 (C) and Laminins (D).

**Supplementary Figure 9.**
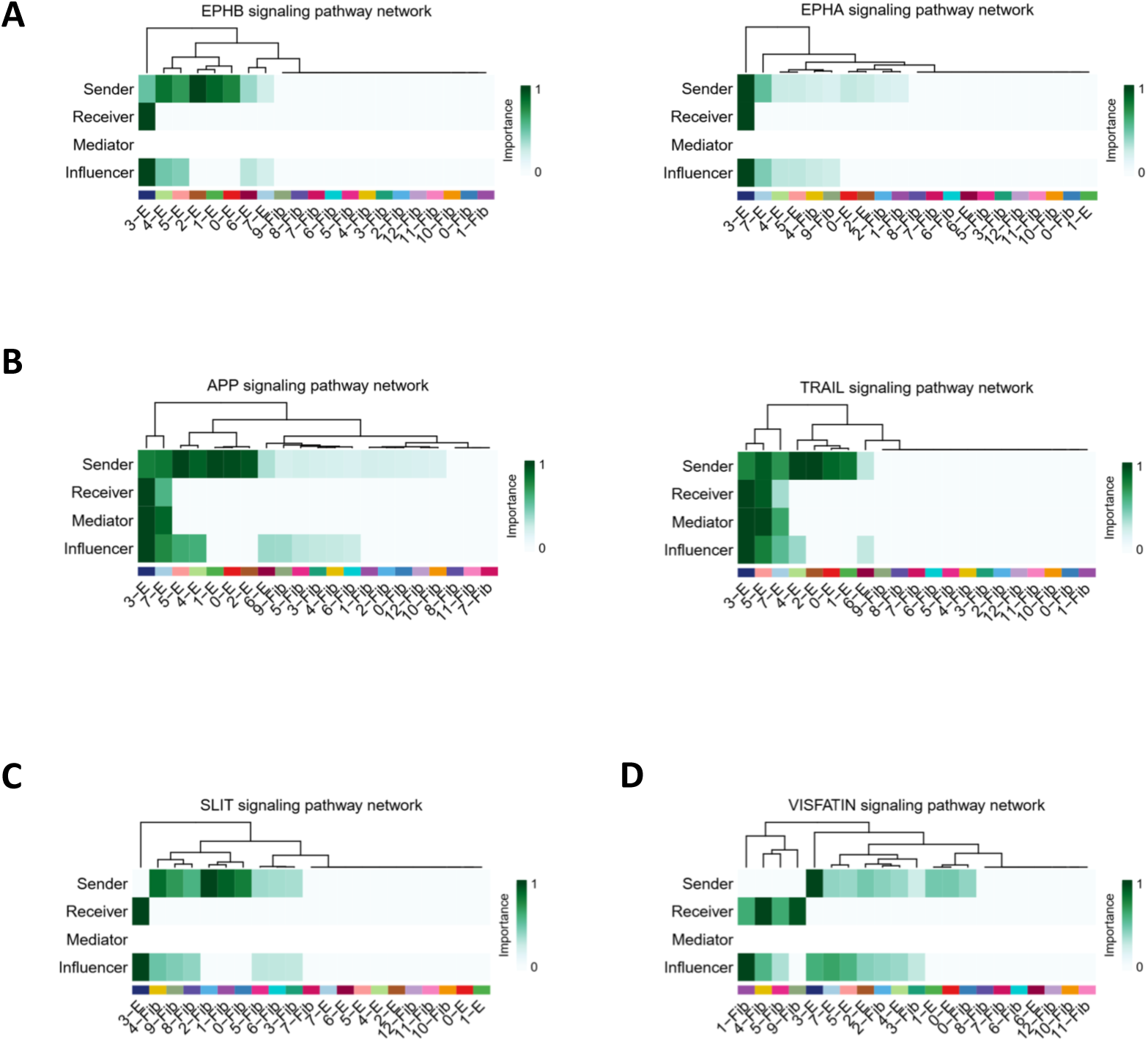
Signalling specific for epithelial Metabolic cluster 3. (**A-C**) Relative contribution of each cell cluster in the signalling primarily received by Cluster 3: ephrin signalling (**A**), pathways involved in TNF signalling (**B**), incoming SLIT signalling from fibroblasts (**C**) and (**D**) outgoing VESFATIN signalling to fibroblasts.

**Supplementary Figure 10.**
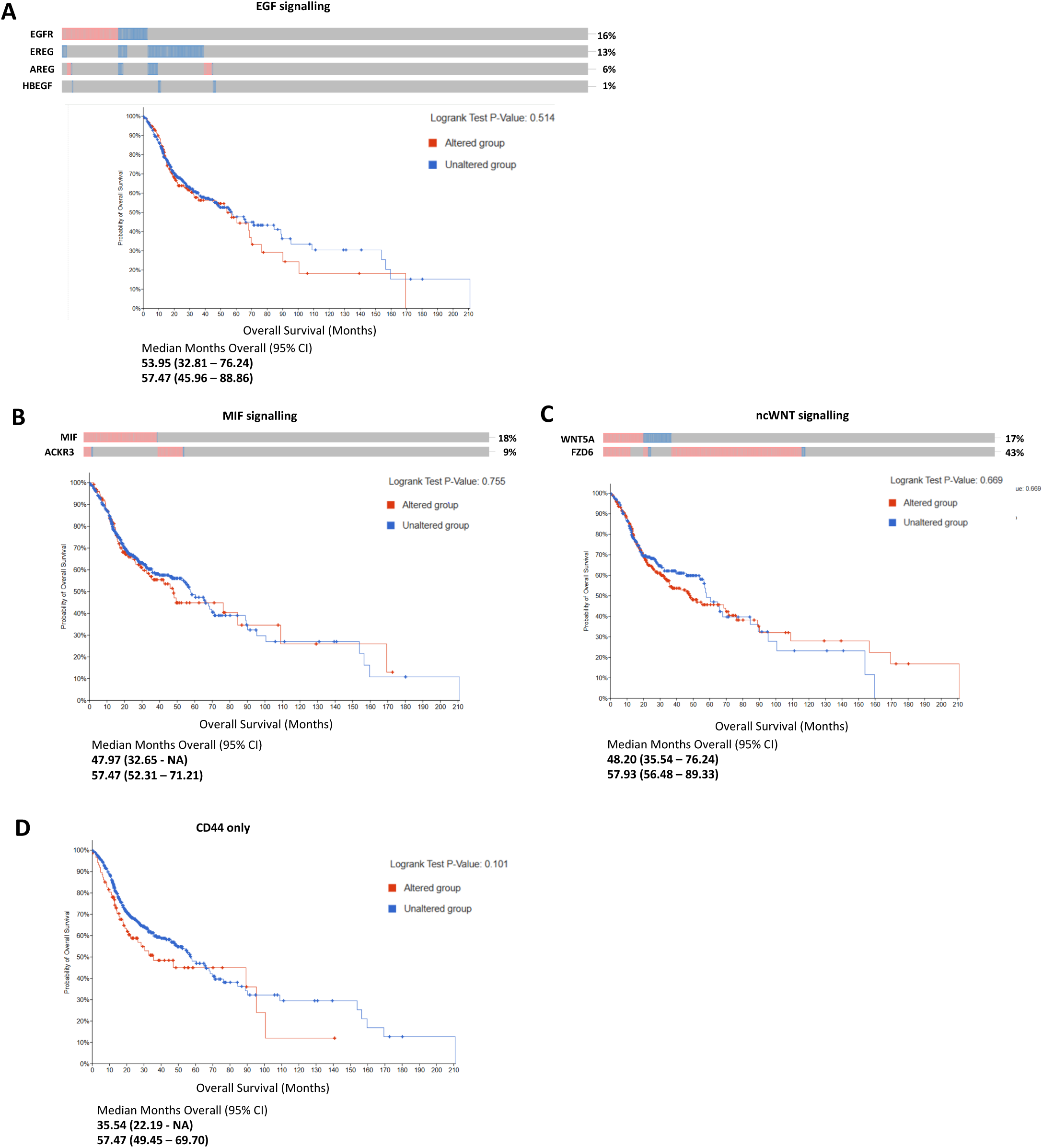
Clinical relevance of the signalling pathways identified in the model. (**A-C**) Overall survival curves for 523 HNSCC patients with data available in TCGA PanCancer cohort with or without upregulation of genes (z-score ≥2.0, relative to normal samples) encoding for the HNSCC-related paracrine signalling pathways: EGF (**A**), MIF (**B**) and ncWNT (**C**). The graph summary shows the difference in median months of overall survival between altered and unaltered groups. The frequency of each gene overexpression in the HNSCC cohort is shown above the graphs. **(D)** Most of the matrix-derived signals are received by epithelial cells via CD44 as shown in Suppl. Fig. 8. Overexpression of these ligand/receptor pairs is associated with lower survival of HNSCC patients (Fig. 7C-D). The Kaplan-Meier curve presented here shows that overexpression of the CD44 receptor alone does not significantly alter patients’ survival.

**Supplementary Table 1.**
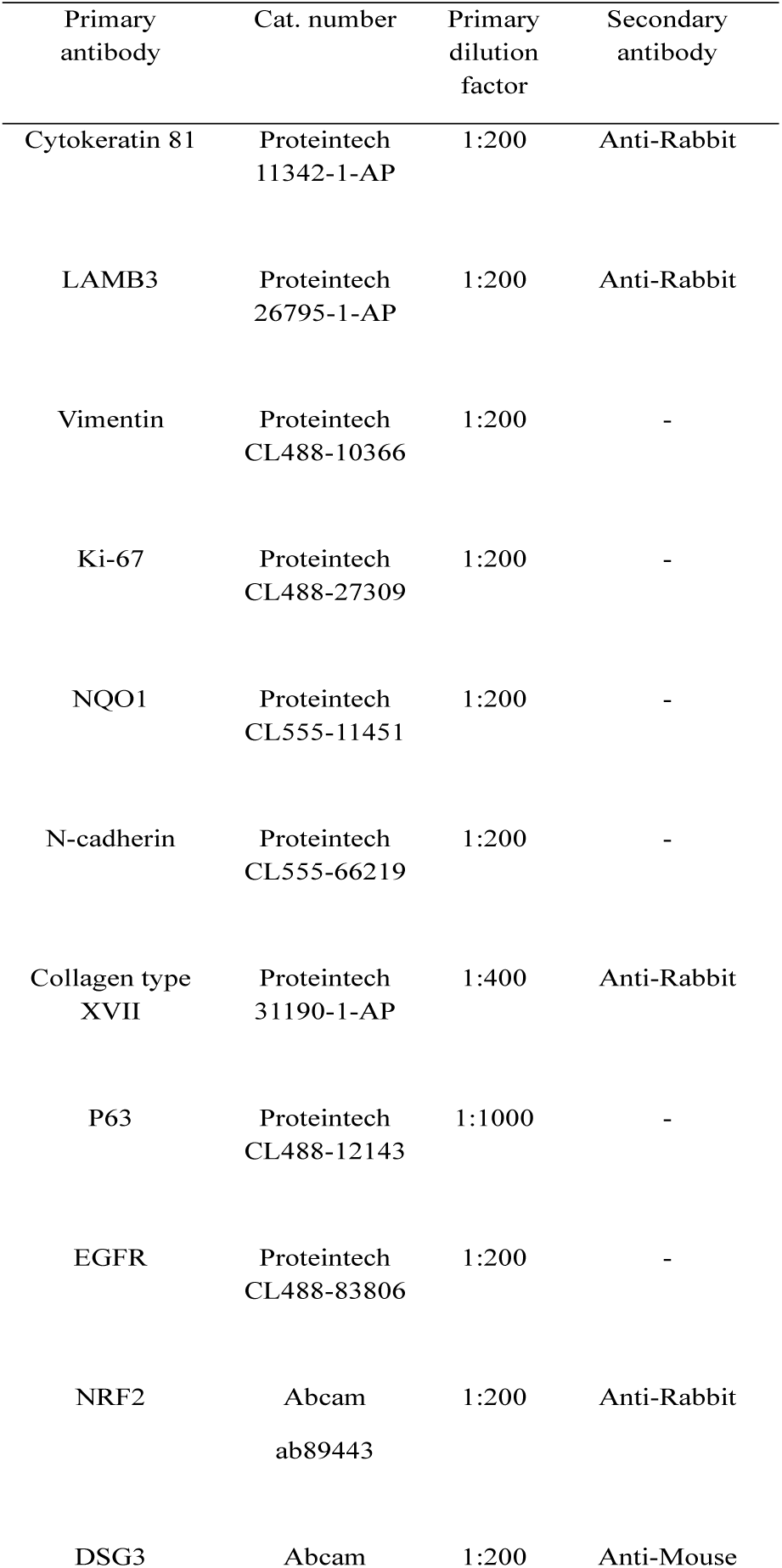
Primary antibodies used in IF staining of organotypic cultures.

| Primary antibody | Cat. number | Primary dilution factor | Secondary antibody |
| --- | --- | --- | --- |
| Cytokeratin 81 | Proteintech<br>11342-1-AP | 1:200 | Anti-Rabbit |
| LAMB3 | Proteintech<br>26795-1-AP | 1:200 | Anti-Rabbit |
| Vimentin | Proteintech<br>CL488-10366 | 1:200 | - |
| Ki-67 | Proteintech<br>CL488-27309 | 1:200 | - |
| NQO1 | Proteintech<br>CL555-11451 | 1:200 | - |
| N-cadherin | Proteintech<br>CL555-66219 | 1:200 | - |
| Collagen type XVII | Proteintech<br>31190-1-AP | 1:400 | Anti-Rabbit |
| P63 | Proteintech<br>CL488-12143 | 1:1000 | - |
| EGFR | Proteintech<br>CL488-83806 | 1:200 | - |
| NRF2 | Abcam<br>ab89443 | 1:200 | Anti-Rabbit |
| DSG3 | Abcam | 1:200 | Anti-Mouse |

## References

1. Andl, C. D., Le Bras, G. F., Loomans, H., Kim, A. S., Zhou, L., Zhang, Y., & Andl, T. (2016). Association of TGFβ signaling with the maintenance of a quiescent stem cell niche in human oral mucosa. Histochemistry and Cell Biology, 146(5), 539–555. 10.1007/S00418-016-1473-0

2. Arora, R., Cao, C., Kumar, M., Sinha, S., Chanda, A., McNeil, R., Samuel, D., Arora, R. K., Matthews, T. W., Chandarana, S., Hart, R., Dort, J. C., Biernaskie, J., Neri, P., Hyrcza, M. D., & Bose, P. (2023). Spatial transcriptomics reveals distinct and conserved tumor core and edge architectures that predict survival and targeted therapy response. Nature Communications, 14(1), 5029. 10.1038/s41467-023-40271-4

3. Bandopadhyay, M., Bulbule, A., Butti, R., Chakraborty, G., Ghorpade, P., Ghosh, P., Gorain, M., Kale, S., Kumar, D., Kumar, S., Totakura, K. V., Roy, G., Sharma, P., Shetti, D., Soundararajan, G., Thorat, D., Tomar, D., Nalukurthi, R., Raja, R., … Kundu, G. C. (2014). Osteopontin as a therapeutic target for cancer. Expert Opinion on Therapeutic Targets, 18(8), 883–895. 10.1517/14728222.2014.925447

4. Bedard, M. C., Chihanga, T., Carlile, A., Jackson, R., Brusadelli, M. G., Lee, D., VonHandorf, A., Rochman, M., Dexheimer, P. J., Chalmers, J., Nuovo, G., Lehn, M., Williams, D. E. J., Kulkarni, A., Carey, M., Jackson, A., Billingsley, C., Tang, A., Zender, C., … Wells, S. I. (2023). Single cell transcriptomic analysis of HPV16-infected epithelium identifies a keratinocyte subpopulation implicated in cancer. Nature Communications, 14(1), 1975. 10.1038/s41467-023-37377-0

5. Belhabib, I., Zaghdoudi, S., Lac, C., Bousquet, C., & Jean, C. (2021). Extracellular Matrices and Cancer-Associated Fibroblasts: Targets for Cancer Diagnosis and Therapy? Cancers, 13(14). 10.3390/CANCERS13143466

6. Bensa, T., Tekkela, S., & Rognoni, E. (2023). Skin fibroblast functional heterogeneity in health and disease. The Journal of Pathology, 260(5), 609–620. 10.1002/path.6159

7. Caetano, A. J., Yianni, V., Volponi, A., Booth, V., D’Agostino, E. M., & Sharpe, P. (2021). Defining human mesenchymal and epithelial heterogeneity in response to oral inflammatory disease. ELife, 10. 10.7554/eLife.62810

8. Cai, H., Liang, J., Jiang, Y., Wang, Z., Li, H., Wang, W., Wang, C., & Hou, J. (2024). KLF7 regulates super-enhancer-driven IGF2BP2 overexpression to promote the progression of head and neck squamous cell carcinoma. Journal of Experimental & Clinical Cancer Research: CR, 43(1). 10.1186/S13046-024-02996-Y

9. Chhabra, Y., & Weeraratna, A. T. (2023). Fibroblasts in cancer: Unity in heterogeneity. Cell, 186(8), 1580–1609. 10.1016/j.cell.2023.03.016

10. Chitty, J. L., & Cox, T. R. (2025). The extracellular matrix in cancer: from understanding to targeting. Trends in Cancer, 11(9), 839–849. 10.1016/j.trecan.2025.05.003

11. Choi, J.-H., Lee, B.-S., Jang, J. Y., Lee, Y. S., Kim, H. J., Roh, J., Shin, Y. S., Woo, H. G., & Kim, C.-H. (2023). Single-cell transcriptome profiling of the stepwise progression of head and neck cancer. Nature Communications, 14(1), 1055. 10.1038/s41467-023-36691-x

12. Constantin, M., Chifiriuc, M. C., Bleotu, C., Vrancianu, C. O., Cristian, R. E., Bertesteanu, S. V., Grigore, R., & Bertesteanu, G. (2024). Molecular pathways and targeted therapies in head and neck cancers pathogenesis. Frontiers in Oncology, 14, 1373821. 10.3389/FONC.2024.1373821

13. Cords, L., Tietscher, S., Anzeneder, T., Langwieder, C., Rees, M., de Souza, N., & Bodenmiller, B. (2023). Cancer-associated fibroblast classification in single-cell and spatial proteomics data. Nature Communications, 14(1), 4294. 10.1038/s41467-023-39762-1

14. Custódio, M., Biddle, A., & Tavassoli, M. (2020). Portrait of a CAF: The story of cancer-associated fibroblasts in head and neck cancer. Oral Oncology, 110. 10.1016/j.oraloncology.2020.104972

15. Deltcheva, E., & Nimmo, R. (2017). RUNX transcription factors at the interface of stem cells and cancer. The Biochemical Journal, 474(11), 1755–1768. 10.1042/BCJ20160632

16. Di Nitto, C., Ravazza, D., Gilardoni, E., Look, T., Sun, M., Prodi, E., Moisoiu, V., Pellegrino, C., Manz, M. G., Puca, E., Weller, M., Weiss, T., Neri, D., & De Luca, R. (2024). An IL-7 fusion protein targeting EDA fibronectin upregulates TCF1 on CD8+ T-cells, preferentially accumulates to neoplastic lesions, and boosts PD-1 blockade. Journal for Immunotherapy of Cancer, 12(8). 10.1136/JITC-2023-008504

17. Eckert, R. L., Adhikary, G., Young, C. A., Jans, R., Crish, J. F., Xu, W., & Rorke, E. A. (2013). AP1 transcription factors in epidermal differentiation and skin cancer. Journal of Skin Cancer, 2013, 1–9. 10.1155/2013/537028

18. Gao, J., Aksoy, B. A., Dogrusoz, U., Dresdner, G., Gross, B., Sumer, S. O., Sun, Y., Jacobsen, A., Sinha, R., Larsson, E., Cerami, E., Sander, C., & Schultz, N. (2013). Integrative analysis of complex cancer genomics and clinical profiles using the cBioPortal. Science Signaling, 6(269). 10.1126/SCISIGNAL.2004088

19. Gari, M. K., Lee, H. J., Inman, D. R., Burkel, B. M., Highland, M. A., Kwon, G. S., Gupta, N., & Ponik, S. M. (2025). Inhibiting fibronectin assembly in the breast tumor microenvironment increases cell death and improves response to doxorubicin. BioRxiv: The Preprint Server for Biology. 10.1101/2025.02.12.637963

20. Gendoo, D. M. A. (2020). Bioinformatics and computational approaches for analyzing patient-derived disease models in cancer research. Computational and Structural Biotechnology Journal, 18, 375–380. 10.1016/J.CSBJ.2020.01.010

21. Germain, P. L., Sonrel, A., & Robinson, M. D. (2020). pipeComp, a general framework for the evaluation of computational pipelines, reveals performant single cell RNA-seq preprocessing tools. Genome Biology, 21(1). 10.1186/S13059-020-02136-7

22. Gleneadie, H. J., Baker, A. H., Batis, N., Bryant, J., Jiang, Y., Clokie, S. J. H., Mehanna, H., Garcia, P., Gendoo, D. M. A., Roberts, S., Burley, M., Molinolo, A. A., Gutkind, J. S., Scheven, B. A., Cooper, P. R., Parish, J. L., Khanim, F. L., & Wiench, M. (2021). The anti-tumour activity of DNA methylation inhibitor 5-aza-2′-deoxycytidine is enhanced by the common analgesic paracetamol through induction of oxidative stress. Cancer Letters, 501, 172–186. 10.1016/J.CANLET.2020.12.029

23. Gougis, P., Bachelard, C. M., Kamal, M., Gan, H. K., Borcoman, E., Torossian, N., Bièche, I., & Le Tourneau, C. (2019). Clinical Development of Molecular Targeted Therapy in Head and Neck Squamous Cell Carcinoma. JNCI Cancer Spectrum, 3(4). 10.1093/JNCICS/PKZ055

24. Groeger, S., & Meyle, J. (2019). Oral Mucosal Epithelial Cells. Frontiers in Immunology, 10(FEB). 10.3389/FIMMU.2019.00208

25. Gronbach, L., Wolff, C., Klinghammer, K., Stellmacher, J., Jurmeister, P., Alexiev, U., Schäfer-Korting, M., Tinhofer, I., Keilholz, U., & Zoschke, C. (2020). A multilayered epithelial mucosa model of head neck squamous cell carcinoma for analysis of tumor-microenvironment interactions and drug development. Biomaterials, 258, 120277. 10.1016/j.biomaterials.2020.120277

26. Haensel, D., Jin, S., Sun, P., Cinco, R., Dragan, M., Nguyen, Q., Cang, Z., Gong, Y., Vu, R., MacLean, A. L., Kessenbrock, K., Gratton, E., Nie, Q., & Dai, X. (2020). Defining Epidermal Basal Cell States during Skin Homeostasis and Wound Healing Using Single-Cell Transcriptomics. Cell Reports, 30(11), 3932–3947.e6. 10.1016/j.celrep.2020.02.091

27. Hewitt, B., Batt, J., Shelton, R. M., Cooper, P. R., Landini, G., Lucas, R. A., Wiench, M., & Milward, M. R. (2022). A 3D Printed Device for In Vitro Generation of Stratified Epithelia at the Air-Liquid Interface. Tissue Engineering. Part C, Methods, 28(11), 599–609. 10.1089/ten.TEC.2022.0130

28. Hiraike, Y., Saito, K., Oguchi, M., Wada, T., Toda, G., Tsutsumi, S., Bando, K., Sagawa, J., Nagano, G., Ohno, H., Kubota, N., Kubota, T., Aburatani, H., Kadowaki, T., Waki, H., Yanagimoto, S., & Yamauchi, T. (2023). NFIA in adipocytes reciprocally regulates mitochondrial and inflammatory gene program to improve glucose homeostasis. Proceedings of the National Academy of Sciences of the United States of America, 120(31), e2308750120. 10.1073/PNAS.2308750120

29. Hurlin, P. J., & Huang, J. (2006). The MAX-interacting transcription factor network. Seminars in Cancer Biology, 16(4), 265–274. 10.1016/j.semcancer.2006.07.009

30. Infante, E., & Etienne-Manneville, S. (2022). Intermediate filaments: Integration of cell mechanical properties during migration. Frontiers in Cell and Developmental Biology, 10, 951816. 10.3389/FCELL.2022.951816/FULL

31. Jin, S., Guerrero-Juarez, C. F., Zhang, L., Chang, I., Ramos, R., Kuan, C.-H., Myung, P., Plikus, M. V, & Nie, Q. (2021). Inference and analysis of cell-cell communication using CellChat. Nature Communications, 12(1), 1088. 10.1038/s41467-021-21246-9

32. Jin, S., Plikus, M. V., & Nie, Q. (2025). CellChat for systematic analysis of cell-cell communication from single-cell transcriptomics. Nature Protocols, 20(1), 180–219. 10.1038/S41596-024-01045-4

33. Johansson, E., & Ueno, H. (2021). Characterization of normal and cancer stem-like cell populations in murine lingual epithelial organoids using single-cell RNA sequencing. Scientific Reports, 11(1), 22329. 10.1038/s41598-021-01783-5

34. Johnson, D. E., Burtness, B., Leemans, C. R., Lui, V. W. Y., Bauman, J. E., & Grandis, J. R. (2020). Head and neck squamous cell carcinoma. Nature Reviews. Disease Primers, 6(1), 92. 10.1038/s41572-020-00224-3

35. Kageyama, R., Ohtsuka, T., & Kobayashi, T. (2007). The Hes gene family: repressors and oscillators that orchestrate embryogenesis. Development (Cambridge, England), 134(7), 1243–1251. 10.1242/DEV.000786

36. Kähler, K. C., Hassel, J. C., Ziemer, M., Rutkowski, P., Meier, F., Flatz, L., Gaudy-Marqueste, C., Zimmer, L., Santinami, M., Russano, F., von Wasielewski, I., Eigentler, T. K., Maio, M., Zalaudek, I., Haferkamp, S., Quaglino, P., Welzel, J., Röcken, C., Enk, A., … Hauschild, A. (2025). Neoadjuvant intralesional targeted immunocytokines (daromun) in stage III melanoma. Annals of Oncology: Official Journal of the European Society for Medical Oncology, 36(10), 1166–1177. 10.1016/J.ANNONC.2025.06.014

37. Kawabe, M., Yang, S., Bolanos, L. G., Takamatsu, S., Flores, S., Castro, P. D., Tsai, T.-Y., Nishikawa-Kaga, A., Frederick, M. J., Sandulache, V., Kadara, H., Myers, J. N., & Osman, A. A. (2025). Targeted suppression of SPP1 inhibits tumor invasion and metastasis in NRF2 hyperactivated cisplatin resistant HNSCC. BioRxiv: The Preprint Server for Biology. 10.1101/2025.05.28.656679

38. Khera, N., Rajkumar, A. S., Abdulkader M, Alkurdi, K., Liu, Z., Ma, H., Waseem, A., & Teh, M.-T. (2023). Identification of multidrug chemoresistant genes in head and neck squamous cell carcinoma cells. In Molecular cancer (Vol. 22, Number 1, p. 146). 10.1186/s12943-023-01846-3

39. Kim, S., Kee, H. J., Kim, D., Jang, J., Jeong, H.-O., Sim, N. S., Selig, M., Ihlow, J., Penter, L., Hwang, T., Choi, D. W.-Y., Lee, K. J., Lee, J., Park, Y. M., Lee, S., & Koh, Y. W. (2025). Multiregional single-cell transcriptomics reveals an association between partial EMT and immunosuppressive states in oral squamous cell carcinoma. IScience, 28(9), 112988. 10.1016/j.isci.2025.112988

40. Kim, S.-Y., Verweij, L. H. G., Lin, L., van Es, J. H., Slack, J., Winkel, C., Candelli, T., Lijnzaad, P., Margaritis, T., Breimer, G. E., Sanders, K., van de Wetering, M., & Clevers, H. (2025). Organoid Modeling of Mouse Anterior Tongue Epithelium Reveals Regional and Cellular Identities. *Advanced Science (Weinheim, Baden-Wurttemberg*, Germany), 12(46), e06738. 10.1002/advs.202506738

41. Kim, Y. J., Lee, H. S., Kim, D., Byun, H. K., Koom, W. S., & Koh, W. G. (2024). Bilayer 3D co-culture platform inducing the differentiation of normal fibroblasts into cancer-associated fibroblast like cells: New in vitro source to obtain cancer-associated fibroblasts. Bioengineering & Translational Medicine, 10(1), e10708. 10.1002/btm2.10708

42. Kohno, Y., Okamoto, T., Ishibe, T., Nagayama, S., Shima, Y., Nishijo, K., Shibata, K. R., Fukiage, K., Otsuka, S., Uejima, D., Araki, N., Naka, N., Nakashima, Y., Aoyama, T., Nakayama, T., Nakamura, T., & Toguchida, J. (2006). Expression of Claudin7 Is Tightly Associated with Epithelial Structures in Synovial Sarcomas and Regulated by an Ets Family Transcription Factor, ELF3. Journal of Biological Chemistry, 281(50), 38941–38950. 10.1074/JBC.M608389200

43. Koster, M. I., & Roop, D. R. (2004). The role of p63 in development and differentiation of the epidermis. Journal of Dermatological Science, 34(1), 3–9. 10.1016/j.jdermsci.2003.10.003

44. Koster, M. I., & Roop, D. R. (2007). Mechanisms regulating epithelial stratification. Annual Review of Cell and Developmental Biology, 23, 93–113. 10.1146/ANNUREV.CELLBIO.23.090506.123357

45. Kumra, H., & Reinhardt, D. P. (2016). Fibronectin-targeted drug delivery in cancer. Advanced Drug Delivery Reviews, 97, 101–110. 10.1016/j.addr.2015.11.014

46. Kürten, C. H. L., Kulkarni, A., Cillo, A. R., Santos, P. M., Roble, A. K., Onkar, S., Reeder, C., Lang, S., Chen, X., Duvvuri, U., Kim, S., Liu, A., Tabib, T., Lafyatis, R., Feng, J., Gao, S.-J., Bruno, T. C., Vignali, D. A. A., Lu, X., … Ferris, R. L. (2021). Investigating immune and non-immune cell interactions in head and neck tumors by single-cell RNA sequencing. Nature Communications, 12(1), 7338. 10.1038/s41467-021-27619-4

47. Lendahl, U., Muhl, L., & Betsholtz, C. (2022). Identification, discrimination and heterogeneity of fibroblasts. Nature Communications, 13(1). 10.1038/S41467-022-30633-9

48. Li, C., Guo, H., Zhai, P., Yan, M., Liu, C., Wang, X., Shi, C., Li, J., Tong, T., Zhang, Z., Ma, H., & Zhang, J. (2024). Spatial and Single-Cell Transcriptomics Reveal a Cancer-Associated Fibroblast Subset in HNSCC That Restricts Infiltration and Antitumor Activity of CD8+ T Cells. Cancer Research, 84(2), 258–275. 10.1158/0008-5472.CAN-23-1448

49. Li, W., Liu, M., Su, Y., Zhou, X., Liu, Y., & Zhang, X. (2015). The Janus-faced roles of Krüppel-like factor 4 in oral squamous cell carcinoma cells. Oncotarget, 6(42), 44480. 10.18632/ONCOTARGET.6256

50. Liu, Z., Mao, X., Xie, Y., Yan, Y., Wang, X., Mi, J., Yuan, H., Zhang, J., Huang, C., Chen, J., Jili, M., Huang, S., Zhang, Q., Wang, F., Mo, Z., & Yang, R. (2025). Single-cell RNA sequencing reveals a fibroblast gene signature that promotes T-cell infiltration in muscle-invasive bladder cancer. Communications Biology, 8(1). 10.1038/s42003-025-08094-9

51. Luecken, M. D., & Theis, F. J. (2019). Current best practices in single-cell RNA-seq analysis: a tutorial. Molecular Systems Biology, 15(6). 10.15252/MSB.20188746

52. Luk, I. Y., Reehorst, C. M., & Mariadason, J. M. (2018). ELF3, ELF5, EHF and SPDEF Transcription Factors in Tissue Homeostasis and Cancer. Molecules (Basel, Switzerland), 23(9). 10.3390/molecules23092191

53. Macartney, R. A., Das, A., Imaniyyah, A. G., Fricker, A. T. R., Smith, A. M., Fedele, S., Roy, I., Kim, H. W., Lee, D., & Knowles, J. C. (2025). In vitro and ex vivo models of the oral mucosa as platforms for the validation of novel drug delivery systems. Journal of Tissue Engineering, 16, 20417314241313456. 10.1177/20417314241313458

54. Magnúsdóttir, E., Kalachikov, S., Mizukoshi, K., Savitsky, D., Ishida-Yamamoto, A., Panteleyev, A. A., & Calame, K. (2007). Epidermal terminal differentiation depends on B lymphocyte-induced maturation protein-1. Proceedings of the National Academy of Sciences of the United States of America, 104(38), 14988–14993. 10.1073/PNAS.0707323104

55. Mateo, J., Berlin, J., De Bono, J. S., Cohen, R. B., Keedy, V., Mugundu, G., Zhang, L., Abbattista, A., Davis, C., Gallo Stampino, C., & Borghaei, H. (2014). A first-in-human study of the anti-α5β1 integrin monoclonal antibody PF-04605412 administered intravenously to patients with advanced solid tumors. Cancer Chemotherapy and Pharmacology, 74(5), 1039–1046. 10.1007/S00280-014-2576-8

56. McConnell, B. B., Ghaleb, A. M., Nandan, M. O., & Yang, V. W. (2007). The diverse functions of Krüppel-like factors 4 and 5 in epithelial biology and pathobiology. *BioEssays: News and Reviews in Molecular*, Cellular and Developmental Biology, 29(6), 549–557. 10.1002/BIES.20581

57. McGinnis, C. S., Murrow, L. M., & Gartner, Z. J. (2019). DoubletFinder: Doublet Detection in Single-Cell RNA Sequencing Data Using Artificial Nearest Neighbors. Cell Systems, 8(4), 329–337.e4. 10.1016/j.cels.2019.03.003

58. Moreno, C. S. (2019). SOX4: The unappreciated oncogene. Seminars in Cancer Biology, 67(Pt 1), 57–64. 10.1016/j.semcancer.2019.08.027

59. Moses, M. A., George, A. L., Sakakibara, N., Mahmood, K., Ponnamperuma, R. M., King, K. E., & Weinberg, W. C. (2019). Molecular Mechanisms of p63-Mediated Squamous Cancer Pathogenesis. International Journal of Molecular Sciences, 20(14). 10.3390/IJMS20143590

60. Mou, T., Zhu, H., Jiang, Y., Xu, X., Cai, L., Zhong, Y., Luo, J., & Zhang, Z. (2023). Heterogeneity of cancer-associated fibroblasts in head and neck squamous cell carcinoma. Translational Oncology, 35, 101717. 10.1016/j.tranon.2023.101717

61. Nieto, M. A., Huang, R. Y. Y. J., Jackson, R. A. A., & Thiery, J. P. P. (2016). EMT: 2016. Cell, 166(1), 21–45. 10.1016/j.cell.2016.06.028

62. Overmiller, A. M., Uchiyama, A., Hope, E. D., Nayak, S., O’Neill, C. G., Hasneen, K., Chen, Y. W., Naz, F., Dell’Orso, S., Brooks, S. R., Jiang, K., & Morasso, M. I. (2024). Reprogramming of epidermal keratinocytes by PITX1 transforms the cutaneous cellular landscape and promotes wound healing. JCI Insight, 9(24). 10.1172/JCI.INSIGHT.182844

63. Parri, M., & Chiarugi, P. (2010). Rac and Rho GTPases in cancer cell motility control. Cell Communication and Signaling: CCS, 8, 23. 10.1186/1478-811X-8-23

64. Passi, A., Vigetti, D., Buraschi, S., & Iozzo, R. V. (2019). Dissecting the role of hyaluronan synthases in the tumor microenvironment. The FEBS Journal, 286(15), 2937–2949. 10.1111/FEBS.14847

65. Pienimäki, J. P., Rilla, K., Fülöp, C., Sironen, R. K., Karvinen, S., Pasonen, S., Lammi, M. J., Tammi, R., Hascall, V. C., & Tammi, M. I. (2001). Epidermal Growth Factor Activates Hyaluronan Synthase 2 in Epidermal Keratinocytes and Increases Pericellular and Intracellular Hyaluronan. Journal of Biological Chemistry, 276(23), 20428–20435. 10.1074/jbc.M007601200

66. Presland, R. B., & Dale, B. A. (2000). Epithelial structural proteins of the skin and oral cavity: function in health and disease. Critical Reviews in Oral Biology and Medicine: An Official Publication of the American Association of Oral Biologists, 11(4), 383–408. 10.1177/10454411000110040101

67. Prince, M. E., Sivanandan, R., Kaczorowski, A., Wolf, G. T., Kaplan, M. J., Dalerba, P., Weissman, I. L., Clarke, M. F., & Ailles, L. E. (2007). Identification of a subpopulation of cells with cancer stem cell properties in head and neck squamous cell carcinoma. Proceedings of the National Academy of Sciences of the United States of America, 104(3), 973–978. 10.1073/pnas.0610117104

68. Puram, S. V, Tirosh, I., Parikh, A. S., Patel, A. P., Yizhak, K., Gillespie, S., Rodman, C., Luo, C. L., Mroz, E. A., Emerick, K. S., Deschler, D. G., Varvares, M. A., Mylvaganam, R., Rozenblatt-Rosen, O., Rocco, J. W., Faquin, W. C., Lin, D. T., Regev, A., & Bernstein, B. E. (2017). Single-Cell Transcriptomic Analysis of Primary and Metastatic Tumor Ecosystems in Head and Neck Cancer. Cell, 171(7), 1611–1624.e24. 10.1016/j.cell.2017.10.044

69. Quah, H. S., Cao, E. Y., Suteja, L., Li, C. H., Leong, H. S., Chong, F. T., Gupta, S., Arcinas, C., Ouyang, J. F., Ang, V., Celhar, T., Zhao, Y., Tay, H. C., Chan, J., Takahashi, T., Tan, D. S. W., Biswas, S. K., Rackham, O. J. L., & Iyer, N. G. (2023). Single cell analysis in head and neck cancer reveals potential immune evasion mechanisms during early metastasis. Nature Communications, 14(1), 1680. 10.1038/s41467-023-37379-y

70. Rivera, C., & Venegas, B. (2014). Histological and molecular aspects of oral squamous cell carcinoma (Review). Oncology Letters, 8(1), 7–11. 10.3892/OL.2014.2103

71. Rodrigues, J., Heinrich, M. A., Teixeira, L. M., & Prakash, J. (2021). 3D In Vitro Model (R)evolution: Unveiling Tumor–Stroma Interactions. Trends in Cancer, 7(3), 249–264. 10.1016/j.trecan.2020.10.009

72. Sahai, E., Astsaturov, I., Cukierman, E., DeNardo, D. G., Egeblad, M., Evans, R. M., Fearon, D., Greten, F. R., Hingorani, S. R., Hunter, T., Hynes, R. O., Jain, R. K., Janowitz, T., Jorgensen, C., Kimmelman, A. C., Kolonin, M. G., Maki, R. G., Powers, R. S., Puré, E., … Werb, Z. (2020). A framework for advancing our understanding of cancer-associated fibroblasts. Nature Reviews. Cancer, 20(3), 174–186. 10.1038/S41568-019-0238-1

73. Sherman, B. T., Hao, M., Qiu, J., Jiao, X., Baseler, M. W., Lane, H. C., Imamichi, T., & Chang, W. (2022). DAVID: a web server for functional enrichment analysis and functional annotation of gene lists (2021 update). Nucleic Acids Research, 50(W1), W216–W221. 10.1093/NAR/GKAC194

74. Sim, R. H. Z., Voon, P. J., Cheo, S. W., & Lim, D. W. T. (2025). Novel Therapeutic Strategies for Squamous Cell Carcinoma of the Head and Neck: Beyond EGFR and Checkpoint Blockade. Biomedicines, 13(8). 10.3390/BIOMEDICINES13081972

75. Sporn, M. B., & Liby, K. T. (2012). NRF2 and cancer: the good, the bad and the importance of context. Nature Reviews. Cancer, 12(8), 564–571. 10.1038/NRC3278

76. Stabell, A. R., Lee, G. E., Jia, Y., Wong, K. N., Wang, S., Ling, J., Nguyen, S. D., Sen, G. L., Nie, Q., & Atwood, S. X. (2023). Single-cell transcriptomics of human-skin-equivalent organoids. Cell Reports, 42(5), 112511. 10.1016/j.celrep.2023.112511

77. Stolte, K. N., Pelz, C., Yapto, C. V., Raguse, J. D., Dommisch, H., & Danker, K. (2020). IL-1β strengthens the physical barrier in gingival epithelial cells. Tissue Barriers, 8(3). 10.1080/21688370.2020.1804249;ISSUE:ISSUE:DOI

78. Strating, E., Verhagen, M. P., Wensink, E., Dünnebach, E., Wijler, L., Aranguren, I., De la Cruz, A. S., Peters, N. A., Hageman, J. H., van der Net, M. M. C., van Schelven, S., Laoukili, J., Fodde, R., Roodhart, J., Nierkens, S., Snippert, H., Gloerich, M., Rinkes, I. B., Elias, S. G., & Kranenburg, O. (2023). Co-cultures of colon cancer cells and cancer-associated fibroblasts recapitulate the aggressive features of mesenchymal-like colon cancer. Frontiers in Immunology, 14, 1053920. 10.3389/fimmu.2023.1053920

79. Sung, H., Ferlay, J., Siegel, R. L., Laversanne, M., Soerjomataram, I., Jemal, A., & Bray, F. (2021). Global Cancer Statistics 2020: GLOBOCAN Estimates of Incidence and Mortality Worldwide for 36 Cancers in 185 Countries. CA: A Cancer Journal for Clinicians, 71(3), 209–249. 10.3322/caac.21660

80. Tanimizu, N., & Mitaka, T. (2013). Role of grainyhead-like 2 in the formation of functional tight junctions. Tissue Barriers, 1(1), e23495. 10.4161/TISB.23495

81. Trapnell, C., Cacchiarelli, D., Grimsby, J., Pokharel, P., Li, S., Morse, M., Lennon, N. J., Livak, K. J., Mikkelsen, T. S., & Rinn, J. L. (2014). The dynamics and regulators of cell fate decisions are revealed by pseudotemporal ordering of single cells. Nature Biotechnology, 32(4), 381–386. 10.1038/NBT.2859

82. Um, J. H., Zheng, Y., Mao, Q., Nam, C., Zhao, H., Koh, Y. W., Shin, S.-J., Park, Y. M., & Lin, D.-C. (2024). Genomic and single-cell characterization of patient-derived tumor organoid models of head and neck squamous cell carcinoma. BioRxiv: The Preprint Server for Biology. 10.1101/2024.06.28.601068

83. van der Veeken, J., Oliveira, S., Schiffelers, R., Storm, G., van Bergen En Henegouwen, P., & Roovers, R. (2009). Crosstalk between epidermal growth factor receptor– and insulin-like growth factor-1 receptor signaling: implications for cancer therapy. Current Cancer Drug Targets, 9(6), 748–760. 10.2174/156800909789271495

84. Wakamori, S., Taguchi, K., Nakayama, Y., Ohkoshi, A., Sporn, M. B., Ogawa, T., Katori, Y., & Yamamoto, M. (2022). Nrf2 protects against radiation-induced oral mucositis via antioxidation and keratin layer thickening. Free Radical Biology and Medicine, 188, 206–220. 10.1016/j.freeradbiomed.2022.06.239

85. Wang, C. Y., Yu, G. T., Gao, C., Chen, J., Li, Q. L., Zhang, L., Wu, M., Sun, Z. J., & Li, L. Y. (2021). Genome-Wide Enhancer Analysis Reveals the Role of AP-1 Transcription Factor in Head and Neck Squamous Cell Carcinoma. Frontiers in Molecular Biosciences, 8. 10.3389/FMOLB.2021.701531/PDF

86. Wang, S. J., & Bourguignon, L. Y. W. (2006). Hyaluronan and the interaction between CD44 and epidermal growth factor receptor in oncogenic signaling and chemotherapy resistance in head and neck cancer. Archives of Otolaryngology--Head & Neck Surgery, 132(7), 771–778. 10.1001/ARCHOTOL.132.7.771

87. Wang, Yinfang, Wan, X., Hao, Y., Zhao, Y., Du, L., Huang, Y., Liu, Z., Wang, Ying, Wang, N., & Zhang, P. (2019). NR6A1 regulates lipid metabolism through mammalian target of rapamycin complex 1 in HepG2 cells. Cell Communication and Signaling: CCS, 17(1). 10.1186/S12964-019-0389-4

88. Xiong, M., Hu, J. J., Yao, M. L., Song, T. T., Zhao, L., Mou, B. Q., Qian, Y. X., Zheng, M. J., Dong, Y. J., Wang, H. Y., Zou, J., & Yang, H. (2024). Single-cell sequencing of head and neck carcinoma: Transcriptional landscape and prognostic model based on malignant epithelial cell features. FASEB Journal: Official Publication of the Federation of American Societies for Experimental Biology, 38(1). 10.1096/FJ.202301287RR

89. Yang, D., Liu, J., Qian, H., & Zhuang, Q. (2023). Cancer-associated fibroblasts: from basic science to anticancer therapy. Experimental & Molecular Medicine, 55(7), 1322–1332. 10.1038/s12276-023-01013-0

90. Yang, S., Feng, T., & Li, H. (2021). KLF5, a Novel Therapeutic Target in Squamous Cell Carcinoma. DNA and Cell Biology, 40(12), 1503–1512. 10.1089/DNA.2021.0674

91. Zappia, L., & Oshlack, A. (2018). Clustering trees: a visualization for evaluating clusterings at multiple resolutions. GigaScience, 7(7). 10.1093/GIGASCIENCE/GIY083

92. Zhang, W., Ge, Y., Cheng, Q., Zhang, Q., Fang, L., & Zheng, J. (2018). Decorin is a pivotal effector in the extracellular matrix and tumour microenvironment. Oncotarget, 9(4), 5480–5491. 10.18632/ONCOTARGET.23869

93. Zheng, G. X. Y., Terry, J. M., Belgrader, P., Ryvkin, P., Bent, Z. W., Wilson, R., Ziraldo, S. B., Wheeler, T. D., McDermott, G. P., Zhu, J., Gregory, M. T., Shuga, J., Montesclaros, L., Underwood, J. G., Masquelier, D. A., Nishimura, S. Y., Schnall-Levin, M., Wyatt, P. W., Hindson, C. M., … Bielas, J. H. (2017). Massively parallel digital transcriptional profiling of single cells. Nature Communications 2017 8:1, 8(1), 14049-. 10.1038/ncomms14049

